# FX-Cell: Quantitative cell release from fixed plant tissues for single-cell genomics

**DOI:** 10.1101/2021.10.11.463960

**Authors:** D. Blaine Marchant, Brad Nelms, Virginia Walbot

## Abstract

Single-cell RNA-sequencing (scRNA-seq) can provide invaluable insight into cell development, cell type identification, and plant evolution. However, the resilience of the cell wall makes it difficult to dissociate plant tissues and release individual cells for single-cell analysis. Here, we show that plant organs can be rapidly and quantitatively dissociated into cells if fixed prior to enzymatic digestion. Fixation enables digestion at high temperatures at which enzymatic activity is optimal and stabilizes the plant cell cytoplasm, rendering cells resistant to mechanical shear force while maintaining high quality RNA. This protocol, FX-Cell, releases four to ten-fold more recoverable cells than optimized protoplasting methods applied to maize anthers or root tips with no cell type biases and can be readily applied to a variety of plant taxa and tissues with no optimization. FX-Cell and scRNA-seq analysis were applied to maize anthers for which 95% of the cells were dispersed and provided suitable scRNA-seq data for the identification of anther cell types with marker genes and well-understood biological functions, including rare meiocytes (∼1% anther cells). In addition, the scRNA-seq data provided putative marker genes and gene ontology information for the identification of unknown cell types. FX-Cell also preserves the morphology of the isolated cells, permitting cell type identification without staining. Ultimately, FX-Cell can be applied to a range of plant species and tissues with minimal to no optimization paving the way for plant scRNA-seq analyses in non-model taxa and tissues.

## INTRODUCTION

The cell holds the genetic blueprint of an organism, yet neighboring cells can differ dramatically in morphology and function. Understanding the gene expression patterns that lead to these differences can provide profound insight into the role, developmental trajectory, and evolution of cell types, tissues, and even organisms. Single-cell RNA sequencing (scRNA-seq) has catalyzed our understanding of animal cells leading to major breakthroughs in cell biology (Han *et al*., 2020), medicine (Lim *et al*., 2020; Paik *et al*., 2020), and evolution (Kanton *et al*., 2019); however, the usage of scRNA-seq in plants has been hampered largely by the presence of the cell wall, which complicates the separation and isolation of single cells (Seyfferth *et al*., 2021).

Plant biologists have largely overcome this hurdle by enzymatically digesting, or protoplasting, the wall of living plant cells (Nelms and Walbot, 2019; Zhang *et al*., 2019; Liu *et al*., 2021; Lopez-Anido *et al*., 2021; Denyer *et al*., 2019). Lacking the cell wall, protoplasts rely on the turgor pressure against the cell membrane for stability making them highly susceptible to bursting via mechanical force or osmotic stress. Generally, protoplasting has been rate-limiting in implementing scRNA-seq as the needed enzymes, enzyme concentrations, digestion time, and digestion conditions vary depending on the species and tissue under investigation. Inadequate protoplasting can result in cell type biases, cell clumps, cell debris, mRNA leakage, or cell lysis, all of which will interfere with the downstream processing needed for scRNA-seq. Even with an optimized protocol, protoplasts are extremely fragile and can have ectopic expression patterns as a result of the lengthy digestion treatment (Denyer *et al*., 2019).

In response to these limitations, one solution has been implementing single nucleus RNA-sequencing (snRNA-seq) in which plant cells are lysed to release the intact nuclei (Conde *et al*., 2021; Sunaga-Franze *et al*., 2021). The nuclei are then isolated and the total nuclear RNA can be reverse transcribed and sequenced. Although snRNA-seq has the benefit of avoiding cell protoplast preparation, nuclear RNA rather than mainly cytoplasmic mRNA is sequenced. As a result, there is significantly less RNA per cell, less sensitive detection of rare transcripts, and an inability to detect distinct isoforms; importantly, nuclear mRNA does not capture the dynamics of translatable mRNAs, which accumulate in the cytoplasm and vary in abundance there over time among different cell types (Sunaga-Franze *et al*., 2021; Thrupp *et al*., 2020).

While recent plant single-cell RNA-seq analyses have begun to diversify in terms of taxa and tissues (Satterlee *et al*., 2020; Nelms and Walbot, 2019; Nelms and Walbot, 2022; Xu *et al*., 2021; Bezrutczyk *et al*., 2021; Bai *et al*., 2022; Zhang *et al*., 2021), the majority of plant scRNA-seq studies have focused on the root tip of *Arabidopsis thaliana* as it has relatively few cell types, established protoplast protocols, and numerous cell type marker genes (Denyer *et al*., 2019; Jean-Baptiste *et al*., 2019; Ryu *et al*., 2019; Zhang *et al*., 2019; Shulse *et al*., 2019). Even in such a well-studied system, these analyses have been instrumental in establishing and refining the spatial and temporal development of the different cell types (Denyer *et al*., 2019; Zhang *et al*., 2019), identifying new marker genes for rare cell types (Denyer *et al*., 2019), and discovering the genetic basis for mutant phenotypes (Ryu *et al*., 2019). Expanding both the taxonomic and tissue diversity of scRNA-seq research in plants promises to address questions related to all realms of basic and applied plant sciences.

Here, we show that cells can be released more efficiently if plant tissues are fixed prior to enzymatic digestion following our novel protocol, FX-Cell. Coagulant fixatives (e.g., Farmer’s Solution, Carnoy’s Solution, Methacarn) stabilize cells by coagulating the protein matrix while removing lipid membranes. We found that fixation followed by cell wall digestion provides two key benefits for cell release: it (i) stabilizes the cell cytoplasm so that cells can withstand harsher shear forces without breaking, and (ii) allows enzymatic digestion to occur at higher temperatures (∼50°C) where the cellulase enzymes are most active (Pardo and Forchiassin, 1999). RNA integrity is maintained by fixation, and high quality RNA can be extracted for later analysis by scRNA-seq. We quantified the cell release of FX-Cell and that of established protoplasting protocols in maize anthers and root tips. We also found that the FX-Cell protocol could readily be applied to a variety of non-model plant systems and maintains cellular morphology after cell wall digestion. This is a critical advancement over previous protoplast-based cell isolation methods as cell morphology is often the sole means of differentiating cell types in taxa and tissues lacking cell type marker genes. To test the genomic suitability of cells released through FX-Cell we performed scRNA-seq on fixed maize anthers. Maize anthers provide an ideal test system for scRNA-seq as the cell type composition of the anther, morphology, and development of the anther cell types are well-documented (Figure 1, A-C) (Kelliher and Walbot, 2011), yet varying degrees of background knowledge (marker genes, biological function, developmental trajectory) exist regarding the genetic activity for each cell type. Meiocytes account for only 1% of the cells in maize anthers, therefore, serve as an exceptional test case for determining if this protocol can be applied to even rare cell types. We demonstrate that FX-Cell can be broadly applied to a variety of taxa and tissues with little to no optimization to provide high-quality scRNA-seq data, thus permitting scalable single-cell research throughout the many study systems of plant biology.

**Figure 1.**
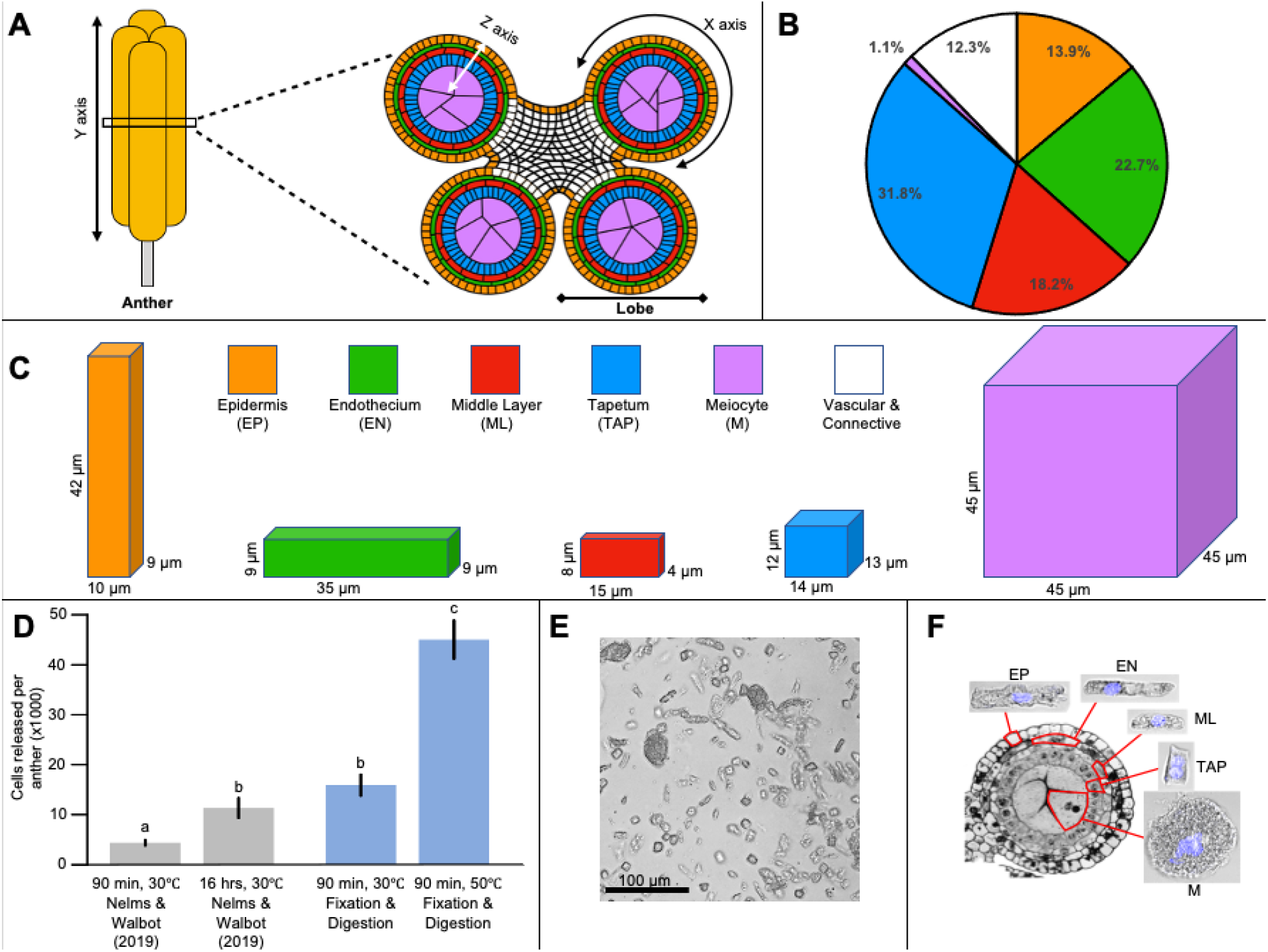
Maize anther anatomy and cellular release with FX-Cell. (A) Transverse section of the maize anther at the beginning of meiosis. (B) Percentage of each cell type per 2.0 mm maize anther based on cell counts from Kelliher & Walbot (2011). (C) Average dimensions for each cell type of the 2.0 mm maize anther from Kelliher & Walbot (2011). Cell types are color coded: orange is epidermis (EP); green is endothecium (EN); red is middle layer (ML); blue is tapetum (TAP); pink is meiocyte (M); white is vascular/connective. (D) Maize anthers were digested for 90 min or 16 h at 30°C following the Nelms and Walbot (2019) protoplasting protocol, and cell release quantified via hemocytometer (grey bars). Fixed maize anthers were digested for 90 min at 30°C and 50°C using the reduced enzyme mix and cell release similarly quantified (blue bars). Fixed maize anther cells after digestion and mechanical dissociation. (F) Transverse section of a maize anther lobe (from Chaubal et al., 2000) with representative images of isolated fixed cells for each anther cell type. Nuclei are shaded blue.

## RESULTS

### Fixation increases cellular release of plant tissues

Cell isolation is perhaps the greatest technical hurdle in scRNA-seq of plant tissues. To determine the possible benefits of fixation on cell isolation, we quantified cell release of fixed plant cells and fresh protoplasts following optimized protocols for both maize anthers and maize root tips. We found that an optimized maize anther protoplast protocol (Nelms and Walbot, 2019) had a mean release of 4,387 cells per anther after 90 min digestion and 11,333 cells per anther when extended to 16 h (Figure 1D). In comparison, if anthers were fixed prior to digestion, 15,900 cells were released within 90 min; this increase in cell release is presumably because the cells were stabilized against mechanical lysis by fixation, while unfixed protoplasts are very fragile. When incubation temperature was increased from 30□ (standard) to 50□ we observed an average release of 45,033 isolated cells (Figure 1D), close to the theoretical number of 50,000 cells in a 2.0 mm maize anther (Kelliher and Walbot, 2011). The standard protoplasting protocol released very few epidermal and endothecial cells, both of which tended to remain clumped and undigested, producing a skewed release favoring tapetal cells, middle layer cells, and meiocytes. When the anthers were fixed then digested at 50□ we did not observe any cell clumps, debris, or undigested material, suggesting that the digestion was complete (Figure 1E). In addition to increasing the cell release efficiency and cell type representation, we found that fixation prior to digestion maintained cells’ natural morphology allowing the potential for cell type identification post-isolation (Figure 1F).

To test the applicability of our fixation-based protocol to another optimized protoplasting protocol of a different maize tissue, we quantified cell release from maize primary root tips after dissociation using: (i) an established maize root tip protoplasting protocol (Ortiz-Ramírez *et al*., 2018); (ii) our fixed-cell method with the enzyme mix from Ortiz-Ramírez et al. (2018); (iii) our fixed-cell method with a reduced enzyme mix. Root tips digested by live tissue protoplasting released 24,667 cells per root tip, similar to what has been reported in the literature (Ortiz-Ramírez *et al*., 2018). In contrast, root tips that were fixed then digested at higher temperatures released approximately four times as many cells using both the protoplasting enzyme mix and our reduced enzyme mix (Figure 2A). Similar to the results we found in anthers, fixed root tips showed little evidence of cell clumps or debris after digestion, suggesting that nearly all cells were released from the tissue (Figure 2B). Protoplasting protocols can be difficult to establish for new tissues. For instance, Ortiz-Ramirez et al. (2018) used a complex protocol to achieve adequate cell release from root tips, including a four-enzyme blend and pretreating live root tissue with L-cysteine. After fixation, we obtained equivalent cell release from roots when using the four-enzyme blend and L-cysteine pretreatment of Ortiz-Ramirez et al. (2018) or using a simpler two-enzyme blend without any treatments (Figure 2A), suggesting the approach might be applied to new tissues with minimal optimization.

**Figure 2.**
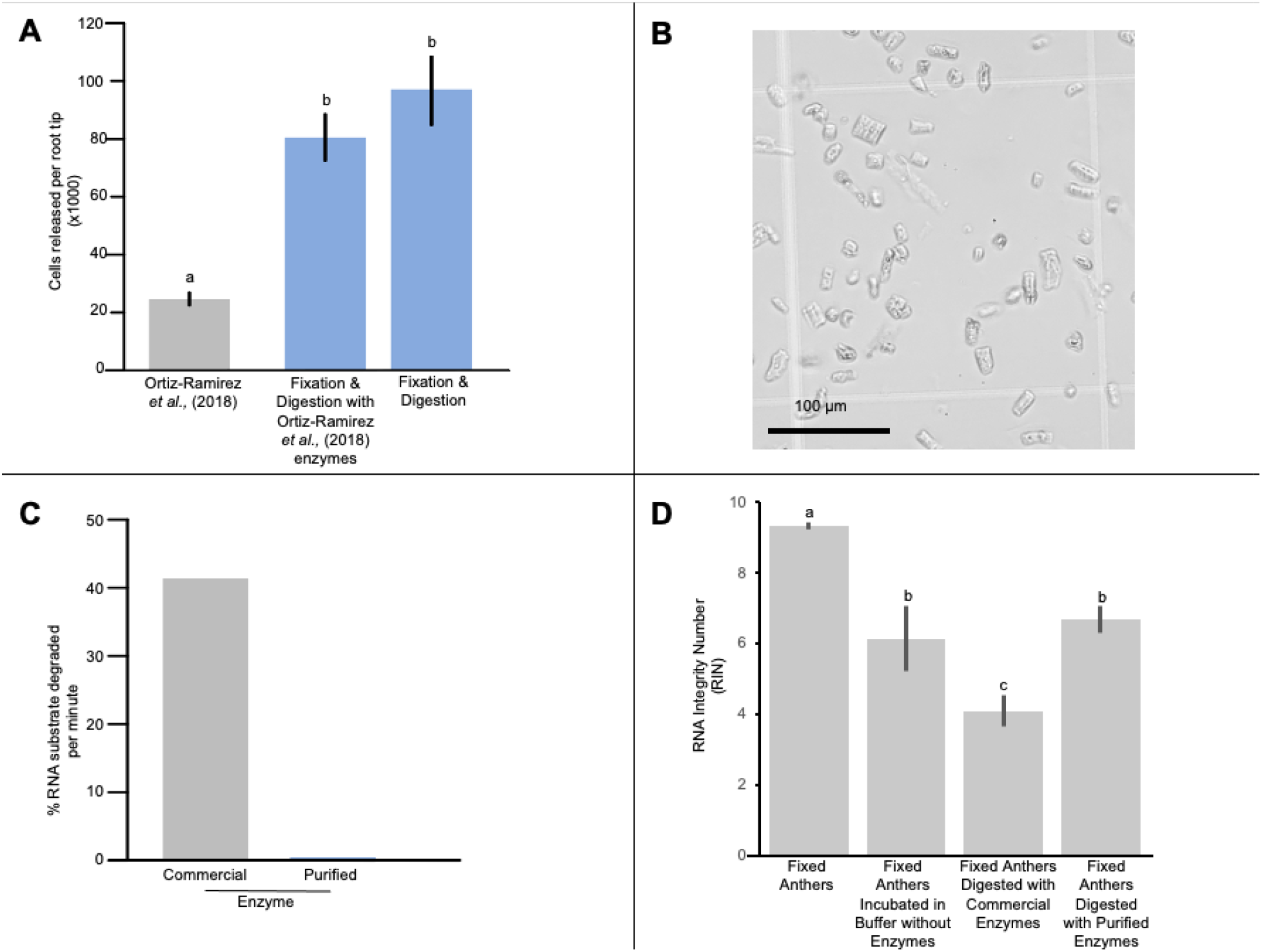
Maize root tip cellular release and RNA integrity of fresh and fixed protoplasting protocols. (A) Cellular release of maize root tips comparing the optimized protoplasting protocol from Ortiz-Ramirez *et al*. (2018) and the fixation-based protocol. Maize root tips were digested following Ortiz-Ramírez *et al*. (2018) and cell release quantified via hemocytometer (grey bars). Maize root tips were fixed then digested at 50°C for 90 min using the Ortiz-Ramírez *et al*. (2018) protoplast enzyme mix and the reduced enzyme mix then cell release was similarly quantified (blue bars). (B) Fixed maize root tip cells after digestion and mechanical dissociation. (C) RNA degradation rate of commercial versus RNase-depleted enzyme mix. (D) RNA quality of fixed maize anthers without any digestion, fixed maize anthers that were digested at 50°C for 90 min in enzyme buffer lacking enzymes, fixed maize anther that were digested at 50°C for 90 min using commercial enzyme mix, and fixed maize anthers that were digested at 50°C for 90 min using RNase-depleted enzyme mix. Different letters denote statistically significant variation (Student’s *t* test, *P* < 0.05) and error bars represent standard error.

To test this hypothesis, we quantified cell release in three additional maize tissues (apical meristem, leaf, young ear) and four non-model plant taxa and tissues (*Amborella trichopoda* leaf, *Nymphaea colorata* leaf, *Capsella bursa-pastoris* leaf and stem) using both our fixation-based, reduced enzyme protocol and a standard protoplasting protocol (Nelms and Walbot, 2019). Although the protoplasting protocol utilized in this experiment was originally optimized for maize anthers, it could serve as a standard starting point for developing a novel protoplasting protocol for new plant tissues or taxa. Cellular morphology was maintained in each fixed sample (Extended Data Fig. 1), allowing obvious differentiation of the varying cell types. Cell release was 10- to 364-fold higher in fixed tissues compared to fresh protoplasts, with the exception of maize leaves in which there were 3.6 as many cells released via fresh protoplasting as by the fixation-based protocol (Extended Data Fig. 2). Presumably, cells with large fluid filled vacuoles, such as maize mesophyll, are very fragile after fixation due to the lack of coagulated proteins; however, testing digestion temperatures, enzyme concentrations, and dissociation methods may surmount even these more difficult cell types. Overall, our fixation-based protocol readily dissociated varying tissues from a breadth of plant species into single cells with no optimization.

### RNase-depletion of enzyme is necessary for maintaining RNA quality

While fixation itself does not affect RNA quality, it removes the cell membrane and makes the internal RNA contents accessible to RNases in solution. This creates a challenge during enzymatic digestion because most cell wall digesting enzymes are complex mixtures that contain substantial RNase activity. We tested several RNase inhibitors, including commercial inhibitors, EDTA, and vanadyl ribonucleoside complexes, but found none that could effectively inhibit the RNase activity in protoplasting enzyme blends. This is partly because many available RNase inhibitors target the RNase A family of enzymes (MacIntosh, 2011), which is only produced in vertebrates. Secreted fungal RNases are primarily of the T1 and T2 families (MacIntosh, 2011).

To surmount this complication, we adapted a column-based method to reduce fungal T1 and T2 RNases by binding them to agarose coupled with guanosine monophosphate (GMP) (Fields *et al*., 1971). We found that cell wall digesting enzymes readily passed through GMP-agarose columns, while the contaminating RNases remained bound. After column depletion, RNase activity was almost completely removed from the enzyme blend (Figure 2C). RNase-depleted enzymes were stable when stored as glycerol stocks for at least a year.

### RNA quality after high temperature digestion

We next tested the effect of the fixed tissue dissociation procedure on RNA quality. RNA isolated from fixed maize anthers had an average RNA Integrity Number (RIN) of 9.3 demonstrating fixation did not cause any significant decrease in RNA quality (Figure 2D). Fixed anthers digested at 50□ in a commercial enzyme blend had a RIN of 4.1 with very noticeable loss of ribosomal RNA. After fixation then digestion with RNAse-depleted enzymes, the RIN was 6.7 demonstrating the fixed tissue dissociation protocol can produce RNA of reasonable quality, although there is a decrease in RNA integrity relative to undigested tissue. When fixed anthers were incubated in enzyme buffer at 50□ without enzymes, we observed a similar RIN of 6.1. Therefore, the decrease in RNA integrity during incubation is not exogenous enzyme-dependent, rather we suspect this degradation is caused by endogenous anther RNases that survive the fixation process. Future improvements of the method may be able to inhibit residual tissue RNases.

### Utilization of FX-Cell for scRNA-seq

Do single cells isolated via FX-Cell have sufficient RNA of high enough quality for scRNA-seq? We prepared four libraries of 96 maize anther cells with FX-Cell (Figure 3A). Two of the libraries were sorted and isolated using a BioSorter (Union Biometrica) and two with a Hana (Namocell). Of the 384 possible single cell samples, 307 had more than 500 UMIs and 200 genes detected after removal of cell-cycle genes. We detected an average of 5,885 UMIs and 2,016 transcribed genes per cell. The dataset was classified into four distinct clusters, two of which were subset and reclustered based on marker gene expression to produce six total clusters (Figure 3B). The total number of UMIs did not vary between the six cell clusters (Extended Data Fig. 3); furthermore, these two independent scRNA-seq experiments using different cell sorting platforms each contributed to the different clusters, indicating that the cell clustering was reproducible between replicates (Extended Data Fig. 3).

**Figure 3.**
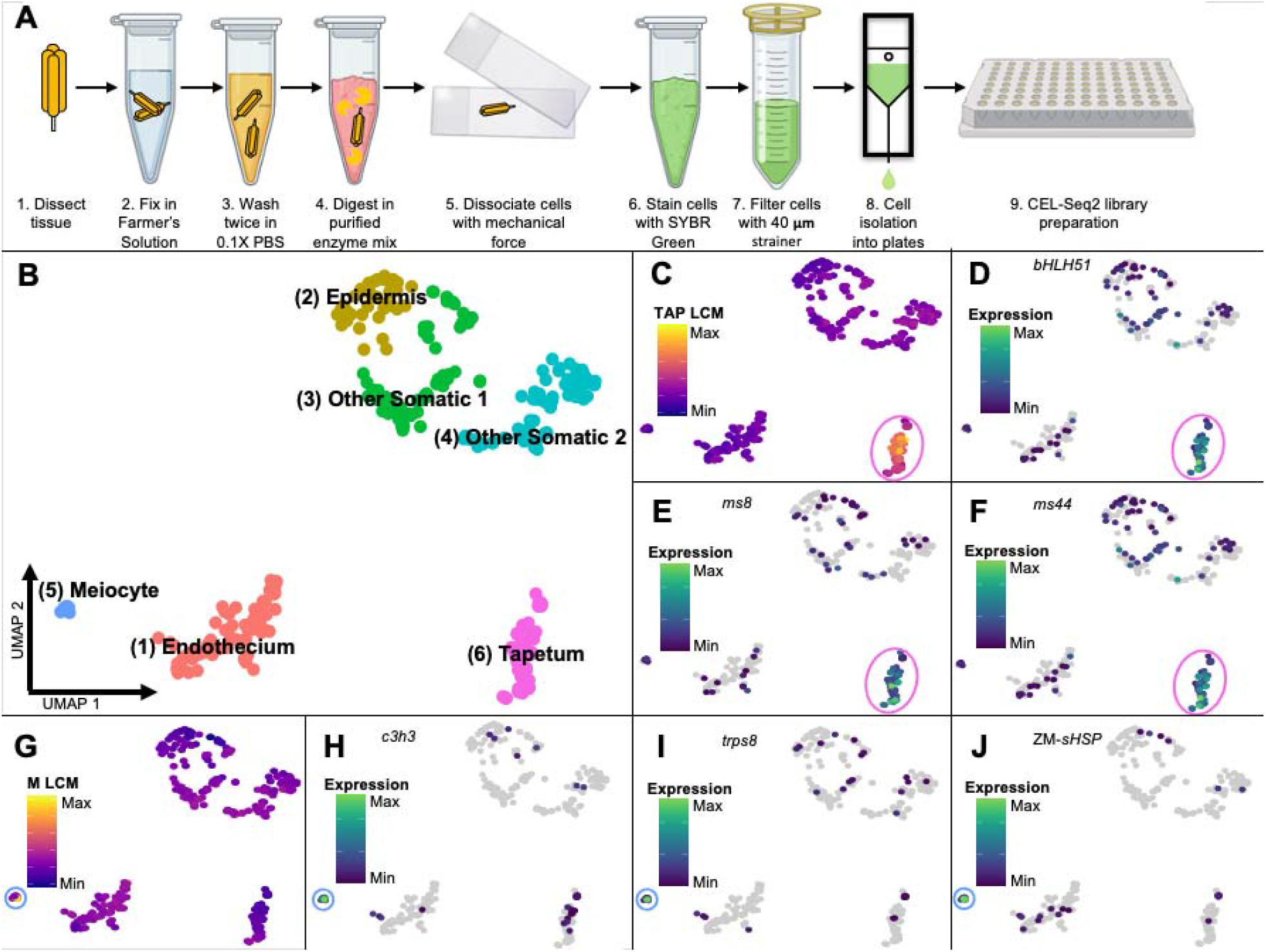
FX-Cell overview and maize anther scRNA-seq of tapetum and meiocyte marker genes. (A) Step-by-step schematic of FX-Cell for scRNA-seq. (B) UMAP clustering of 307 cells from 2.0 mm maize anthers into six distinct clusters. (C) Correlation values of each cell with LCM tapetal data. (D-F) Expression values per maize anther cell of tapetal marker genes mapped onto the UMAP clusters. (G) Correlation values of each cell with LCM meiocyte data. (H-J) Expression values per maize anther cell of meiocyte marker genes mapped onto the UMAP clusters.

We next asked if the FX-Cell scRNA-seq data was sufficient enough to associate the cell clusters with established maize anther cell types based on known marker genes and gene expression of anther cell types isolated by laser capture micro-dissection (LCM) (Zhou *et al*., 2022). We observed a strong correlation between the genes expressed in Cluster 6 and genes expressed in tapetal cells by LCM (Figure 3C; Extended Data Fig. 3). Cluster 6 further expressed several male-sterility genes known to be up-regulated in the tapetum: *basic Helix-Loop-Helix 51* (*bHLH51*), *Male-sterile 8* (*Ms8*), and *Male-sterile 44* (*Ms44*) (Nan *et al*., 2017; Wang *et al*., 2010; Fox *et al*., 2017) (Figures 3, D-F; Extended Data Fig. 4). Genes expressed in Cluster 5 were highly correlated with the LCM meiocyte sample and also had strong expression of genes known to be highly expressed in meiocytes: *Trehalose 6-Phosphate Phosphatase* (*Trps8*), *C3H Transcription Factor 33* (*C3H3*), and a *Small Heat Shock Protein* (*sHSP*) (Nelms and Walbot, 2019; Zhou *et al*., 2022) (Figure 3, G-J; Extended Data Fig. 3, 4). Based on these data, we conclude that Cluster 6 contains tapetal cells and Cluster 5 contains meiocytes.

**Figure 4.**
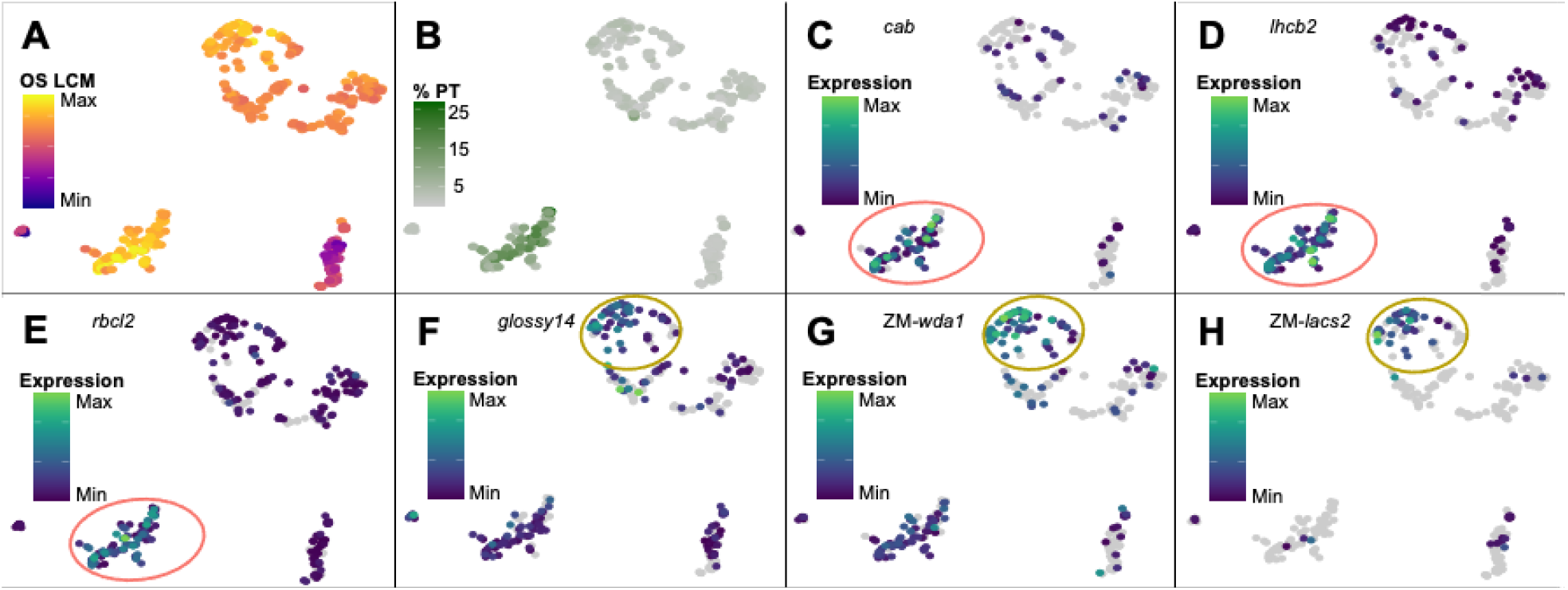
Endothecium and epidermis marker gene expression in maize anther cells. (A) Correlation values of each cell with LCM other somatic cell types (middle layer, endothecium, epidermis) data. (B) Percentage of total UMIs originating from the plastid for each cell. (C-E) Expression values per maize anther cell of putative endothecium marker genes mapped onto the UMAP clusters. (F-H) Expression values per maize anther cell of putative epidermis marker genes mapped onto the UMAP clusters.

The remaining cell clusters all showed low correlation with the LCM tapetal and meiocyte samples but high correlation with the LCM sample data consisting of other somatic cells (epidermis, endothecium, middle layer) (Figure 4; Extended Data Fig. 3). Beyond the tapetum, the maize anther contains multiple different somatic cell types including middle layer, endothecium, epidermis, connective, and vasculature. There is no expression data for these anther cell types and so we attempted to associate the remaining clusters to cells based on knowledge of anther cell biology. Murphy *et al*. (2015) discovered that the endothecium contains chloroplasts unlike the other anther cell layers. We found plastid transcripts were more highly expressed in Cluster 1 than in any other cluster (Figure 4B; Extended Data Fig. 3). Furthermore, transcripts for the photosynthesis-associated genes identified by Murphy *et al*., (2015) and nuclear-encoded chloroplastic proteins (PantherDB Family #21649) (Mi *et al*., 2021) were selectively expressed in Cluster 1 (Figures 4, C-E; Extended Data Fig. 4). Thus, we assign Cluster 1 as endothecium.

The anther epidermis produces cuticular waxes to seal and protect the maize anther from the environment. These waxes are formed by converting C_2_ acetyl-coenzyme A (acetyl-CoA) into C_16_ or C_18_ fatty acids then further converted into fatty acyl-CoAs by long-chain acyl-CoA synthetases (LACS); these are remodeled and extended into C_24_ to C_34_ fatty acids, or very-long-chain fatty acids (VLCFAs) (Zheng *et al*., 2019; Schnurr *et al*., 2004). A number of genes have been found to regulate the production of theses epicuticular waxes in maize, rice, and *Arabidopsis* (Zheng *et al*., 2019; Jung *et al*., 2006; Schnurr *et al*., 2004). We focused on *Glossy14*, the maize homolog of rice *Wax-Deficient Anther1* (*Wda1*), and the maize homolog of *Arabidopsis LACS2* – mutations in these three genes have been shown to result in significantly decreased epicuticular wax load (Zheng *et al*., 2019; Jung *et al*., 2006; Schnurr *et al*., 2004). We found that Cluster 2 had the highest average expression levels and proportion of cells expressing these genes, suggesting Cluster 2 is epidermis (Figures 4, F-H; Extended Data Fig. 4).

The final two clusters were unidentifiable as little is known about the genetic activity of the remaining somatic cell types: the middle layer, connective cells, and vasculature. However, we were able to generate a list of the most specifically expressed genes for each cluster, providing a putative list of marker genes (Figure 5). We were also able to identify modules of co-regulated genes specific to each cluster (Figure 5B). Cluster-specific modules can be analyzed for gene ontology (GO) term enrichment providing insight into the biological processes, cellular component, and molecular function and of each module. For example, Module 31, which was highly up-regulated in the endothecium cluster, is highly enriched for genes relating to photosynthesis and localized in the chloroplast (Figure 5C). Similar analyses can be utilized to verify cluster identification or further narrow down the cell type identity of unknown clusters.

**Figure 5.**
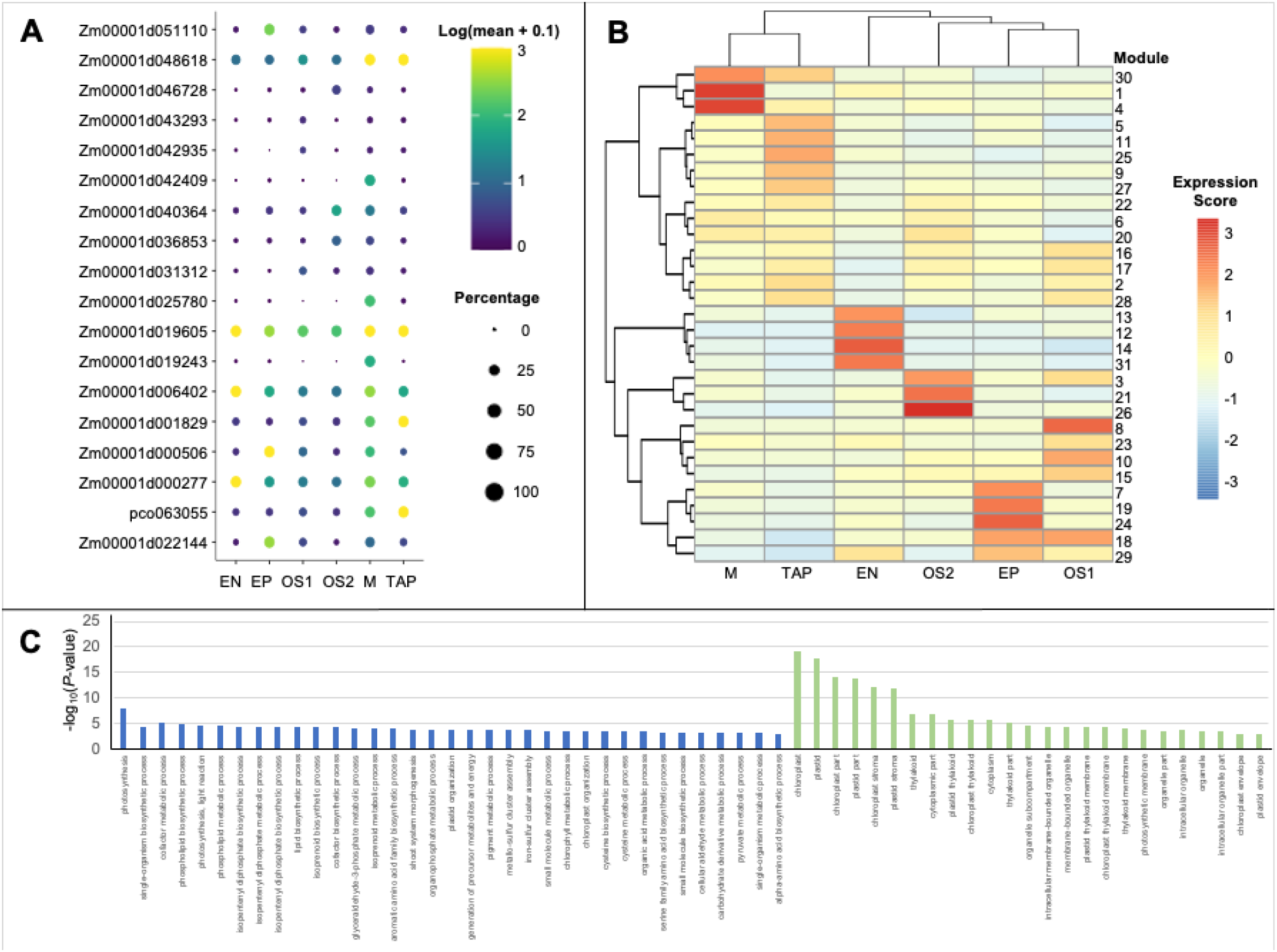
*De novo* marker gene identification and co-regulated gene modules. (A) Top three marker genes for each UMAP cluster with expression and percentage of cells from each cluster expressing the gene. (B) Co-regulated gene modules and expression scores for each UMAP cluster. (C) Gene ontology (GO) enrichment analysis of Module 31 sorted by biological processes (blue bars) and cellular component (green bars). Significantly enriched GO terms were selected based on a false discovery rate (FDR) < 0.05.

## DISCUSSION

Difficulties in the dissociation of tissues and isolation of single cells have restricted plant single-cell RNA-sequencing to only the most researched plant species and tissues. FX-Cell can be readily adapted to an array of plant taxa and tissues spanning well beyond typical model plant species and tissues for single-cell molecular analyses. By incorporating cellular fixation and cell wall digestion enzymes depleted of RNases, we demonstrated that FX-Cell had a significantly higher and more representative cell release than well-established fresh protoplasting protocols in multiple tissues and species while maintaining high-quality RNA with minimal or no additional optimization. Fixation stabilizes cells by coagulating the entire cytosol into a protein matrix making them more resistant to mechanical force and permitting the use of increased digestion temperatures relative to highly fragile and environmentally sensitive fresh protoplasts. FX-Cell is also highly scalable, permitting the isolation and sequencing of a few cells isolated by hand to thousands of cells isolated and dispensed with a cell sorter.

Two technologies for isolating large plant cells in high-throughput applications were identified: the BioSorter (Union Biometrica, Inc.) and Hana (Namocell). These two technologies can readily sort and dispense single fixed plant cells of varying sizes and shapes into plates for library preparation. Although we utilized a modified CEL-Seq2 library preparation protocol for our scRNA-seq analyses, SPLiT-seq (Rosenberg *et al*., 2018), a relatively new and inexpensive way to construct scRNA-seq libraries, requires fixed cells and is well-suited to work smoothly with FX-Cell.

FX-Cell provided high-quality cells for scRNA-seq. We identified tapetal, epidermal, endothecial, and meiocyte cells from fixed maize anthers based on a few known cell type marker genes and the established biology of specific cell types (Figure 3). The notable presence of meiocytes (∼1% of anther cells) in our scRNA-seq dataset validated the ability of FX-Cell to liberate and hence distinguish even rare cell types. The remaining unknown cells likely consist of the middle layer, vascular, and connective cells, but little is known about the expression patterns of these somatic cell types. For example, the function of the middle layer is completely unknown, although its developmental origin and fate are well-established in maize. This ephemeral cell layer differentiates from the secondary parietal cells along with the tapetum early in anther development then undergoes programmed cell death prior to the completion of meiosis. A few male-sterile maize mutants have aberrant middle layer phenotypes, however, the cell layer has been largely understudied relative to the tapetum (Walbot and Egger, 2016). Targeted analysis of this enigmatic cell layer using scRNA-seq could reveal its function and activity in the anther.

It is entirely possible that unknown cell types exist among the vascular and connective tissues or even among the primary four somatic layers of the anther, as demonstrated by Murphy et al., (2015) with the subclassification of the endothecium into the subepidermal endothecium and interendothecium, the endothecial cells adjacent to the connective tissue. In addition, maize tapetal cells asynchronously become binucleate throughout meiosis, suggesting a key developmental transition in this cell type. The substructure of the tapetal cluster may reflect this cellular change or the binucleate tapetal cells could be clustered in the unidentified clusters. Increased sampling and the incorporation of developmental trajectories would heighten the resolution of each cell cluster revealing unknown and unresolved cell types.

The *de novo* identification of the top specific marker genes and co-regulated gene modules for each cluster can help elucidate the identity of unknown scRNA-seq cell clusters. RNA *in situ* hybridization of these putative marker genes could locate these cells within the maize anther, while LCM RNA-seq of the known cell layers could serve as a background reference. GO term enrichment analyses of the co-regulated gene modules can provide critical insight into the function and biology of unknown cell clusters. Coupled with scRNA-seq, high-throughput imaging of each fixed cell before library preparation can categorize cells based on size and shape traits that differ considerably among plant cell types, but these traits are eliminated by protoplasting.

With any new method, it is important to consider potential limitations. The advantage of our method is that it dramatically increases the release of cells from plant tissues. However, there are some contexts where this method has drawbacks. First, some plant cells have large fluid filled vacuoles and are very fragile after fixation; for instance, we found maize leaf mesophyll cells do not hold up well to our method. As a result, other approaches may be better for cells with very high water content. With any new tissue, we recommend first testing this method using commercially available enzymes to see how well the cells of interest are successfully released before committing to RNase-depletion of the enzymes.

Second, we suspect the method will not be compatible with widely used droplet-based technologies such as 10X Genomics. This is because the large size and unusual shape of many plant cells (10 - 100 µm) relative to animal cells (10 - 30 µm) might result in clogging of the microfluidic chips used for droplet-based scRNA-seq with fixed cells. Elongated plant cells become spherical via protoplasting making their overall dimensions more feasible for the microfluidic channels of droplet-based technologies. In addition, we surmise that if a protoplast is too large and blocks the entrance of the microfluidic channel, it will likely lyse via pressure. This prevents clogging of the chip, but also biases the downstream analyses as larger cells will be selectively removed. In contrast, fixed cells will maintain their natural, elongated shapes and are too stable to lyse due to pressure, making the chances of clogging much higher.

Single-cell RNA-seq has revolutionized our understanding of animal cell identification, development, and evolution over the last two decades while scRNA-seq in plants has been slow to develop, largely reflecting the extensive optimization required for dissociating and isolating plant cells. FX-Cell should similarly open the door to such discoveries for plant research regardless of species or tissue.

## METHODS

### Plant growth and anther dissection

*Zea mays* (inbred line W23 *bz2*) individuals were grown under greenhouse conditions in Stanford, CA, USA with 14-h day/10-h night lighting. Daily irrigation and fertilization were maintained for robust growth. Beginning five to six weeks after planting, individual plants were felled ∼20 cm above ground level for anther dissection between 8:00 and 9:00 am. The sacrificed plants were taken to the lab within 10 min where the tassels were dissected out of the stem and leaf whorl. A Leica M60 dissecting scope (Leica Microsystems Inc.) and stage micrometer (Fisher Scientific) were used to isolate 2.0 mm anthers from the upper florets of spikelets along the central spike of the tassel.

### Cellular release

Cell release of fixed and fresh maize anthers was compared in a variety of conditions. Three 2.0 mm anthers were pooled per replicate with five replicates per condition. Fresh anthers were digested at 30□ for 90 min or 16 h in the enzyme mix from Nelms & Walbot (2019). Fixed samples were left in ice-cold Farmer’s solution (3:1 100% ethanol:glacial acetic acid) for two h, washed twice in ice-cold 0.1X phosphate-buffered saline (PBS; Sigma-Aldrich) for five min, then digested at 30□ or 50□ for 90 min in 20 mM MES, pH 5.7 with a 1:10 dilution of RNase-depleted cell wall digesting enzyme stock (enzymes were stored in glycerol stocks, see below; stocks were normalized so that a 1:10 dilution has the same A280 as a 1.25% w/v Cellulase-RS and 0.4% w/v Macerozyme-R10 solution). The cells from the digested, fixed anthers were dissociated via shear force between two microscope slides with thin tape as a spacer. For each replicate, the number of single cells was estimated using a hemocytometer then averaged. Images of the dissociated cells were taken on a Nikon Diaphot inverted microscope with a mounted Nikon D40 camera.

Cell release from maize root tips was compared in three conditions: fresh protoplasting following Ortiz-Ramirez *et al*. 2018, fixation and digestion with the enzyme concentrations from Ortiz-Ramirez *et al*. 2018, or fixation and digestion with our highly reduced enzyme mix. Maize seeds were treated, germinated, and grown following Ortiz-Ramírez *et al*. (2018). Seedling primary roots were cut 5 mm above the tip with a scalpel. Three root tips were pooled per replicate with five replicates per condition. Fresh root tips were pre-treated, washed, digested in enzyme (1.2% Cellulase-RS, 0.36% Pectolyase Y-23, 0.4% Macerozyme-R10, 1.2% Cellulase-R10; Sigma Aldrich; Yakult Pharmaceutical Industry Co.), filtered, and washed. The fresh protoplasts were counted with a hemocytometer. The fixed samples were left in ice-cold Farmer’s solution for two h, washed twice with ice-cold 0.1X PBS for five min then digested at 50□ with the enzyme mix from Ortiz-Ramírez *et al*. (2018) or the enzyme mix from this protocol. Digested tissue was manually disrupted with pipetting, then the number of individual cells counted with a hemocytometer.

We expanded our sampling by comparing the cell release of FX-Cell to a standard protoplasting protocol (Nelms and Walbot, 2019) in three additional maize tissues (apical meristems, young leaves, young ears) and four non-model plant taxa and tissues (leaves from the basal angiosperms, *Amborella trichopoda* and waterlily, *Nymphaea colorata*, leaf and stem tissue from the non-model Brassicaceae *Capsella bursa-pastoris*). For each tissue, comparably sized samples were either fixed in Farmer’s solution, washed twice 0.1X PBS, and digested at 50□in our reduced enzyme mix as previously described or directly digested at 30□ for 90 min in the enzyme mix from Nelms and Walbot (2019). For each replicate, the number of single cells was estimated using a hemocytometer then averaged. Images of the dissociated cells were taken on a Nikon Diaphot inverted microscope with a mounted Nikon D40 camera.

### RNase-depletion of enzymes

RNases present in fungal cell wall digesting enzymes were depleted by passing concentrated enzyme solution through agarose beads coupled with guanosine monophosphate (GMP). GMP beads were prepared using the procedure from Kanaya & Uchida (1981), with modifications: 50 mL suspended ω-aminohexyl–agarose beads (Sigma-Aldrich) were washed three times in water and then three times in 0.1M borax, pH 9.0 (Sigma-Aldrich). Meanwhile, sodium metaperiodate (Chem-Impex) was dissolved in 6 mL water to a final concentration of 0.2 M, and 488 mg guanosine monophosphate was added; the solution was incubated at room temperature (RT) in the dark for 1 h with gentle mixing. The washed agarose beads were resuspended in 0.1 M borax, pH 9.0 to a total volume of 36 mL, then the 6 mL solution containing oxidized GMP was added and the reaction was incubated at RT with gentle mixing for 2-4 h. Finally, 136 mg of solid sodium borohydride (Sigma-Aldrich) was slowly added to the reaction, and the solution was gently mixed at 4□ for 1 h with the cap loosened to allow ventilation. The coupled GMP beads were washed three times each with 0.1 M borax, then water, then 1 M sodium chloride. Washed beads were loaded into a cleaned out Superdex 200 10/300 FPLC column (Cytiva) and stored in 1 M sodium chloride until further use. Remaining beads were stored in a sealed container in 1 M sodium chloride until further use.

For RNase-depletion, the enzymes were resuspended at 10X concentration (12.5% w/v Cellulase-RS and 4% w/v Macerozyme R10) in RNase binding buffer (RBB; 150 mM NaCl, 10 mM citrate, pH 7.0). Four mL of GMP beads were loaded in a Kontes Flex-Column (Kimble Chase) gravity flow column and equilibrated with RBB at 4□, then the enzyme mix was passed through this column. The flow through was collected and then run through the pre-equilibrated GMP-agarose FPLC column at 4□ using a peristaltic pump. Fractions were collected, and those with >0.1 absorbance at A280 were pooled. Pooled enzymes were concentrated using an Amicon Ultra-15 Centrifugal Filter Unit, MWCO 30 kDa (MilliporeSigma) at 4□ until a 1:10 dilution of the enzyme blend had 0.75 absorbance at A280. The Amicon concentrators were made using regenerated cellulase esters and the concentrated enzyme blend was capable of weakening these membranes; for future RNase-depletions, it is recommended to use a centrifugal concentrator with a membrane made from a different material. Concentrated enzymes were mixed 1:1 with glycerol and stored at -20□ until further use. For digestions, enzyme stocks were used at 1/10^th^ the final volume. RNase activity in the RNase-depleted and commercial enzyme mix was quantified using the Ambion RNaseAlert Lab Test Kit (Invitrogen).

### RNA integrity

Anther RNA quality was tested in three conditions of cell preparation: 1) fixed in Farmer’s solution then washed twice in 0.1X PBS then flash frozen, 2) fixed, washed, and digested in commercial enzyme, and 3) fixed, washed, and digested in RNase-depleted enzyme. For each condition, 2.0 mm anthers were isolated from five separate plants with ten anthers pooled per plant. The flash frozen samples were homogenized via bead beating in a 2000 Geno/Grinder (SPEX CertiPrep) with baked 4 mm steel balls. The fixed samples were left in ice-cold Farmer’s solution for two h, washed twice in ice-cold 0.1X PBS for five min, then incubated at 50□ for 90 min with RNase-depleted or commercial enzyme (1.25% w/v Cellulase-RS and 0.4% w/v Macerozyme R10). The RNeasy Plant Mini Kit (Qiagen) was used to extract RNA from samples via the standard protocol. RNA was quality-checked on an Agilent 2100 BioAnalyzer with the RNA 6000 Nano assay (Agilent Technologies). The RNA Integrity Number (RIN) for the five replicates of each condition were averaged and reported alongside the error.

### Fixed cell isolation for scRNA-Seq

Anthers from four individuals of wild-type (W23) maize were dissected out. One of the three anthers per floret was used for imaging on a Nikon Diaphot inverted microscope with a Nikon D40 mounted camera at 10X magnification. The remaining two anthers per floret were fixed in ice-cold Farmers solution for two h, washed twice for 5 min in 0.1X PBS, and then one anther was digested for 90 min at 50□ in the RNase-depleted enzyme mix while the other anther was saved at -20□. Following digestion, shear force was applied to the anther between two microscope slides with thin tape on each end to prevent the anther from being fully crushed. The top microscope slide was slid back and forth 5-10 times and the sample checked under the dissecting scope to ensure separation of the fixed cells. The cells were washed from the slides into 1 mL of cold 0.1X PBS via pipette and stained with SYBR Green I nucleic acid gel stain (Invitrogen) for 20 min. The cells were then filtered through a 100 µm (if bound for the BioSorter) or 40 µm (if bound for the Hana) nylon cell strainer (Corning Inc.) into 50 mL Falcon tubes. The stained cells were then sorted into 384-well plates or 96-well plates, each well containing 0.8 µL Primer Master Mix (0.225% Triton X-100, 1.6 mM dNTP mix, 1.875 uM barcoded oligo[dT] CEL-seq2 primers; Sigma-Aldrich, New England Biolabs) using a BioSorter (Union BioMetrica) or Hana Single Cell Dispenser (Namocell). Following cell sorting, the plates were spun at 400 x *g* then stored at -80□.

### CEL-Seq2 library preparation

Single cell libraries were prepared following the CEL-seq2 protocol(Hashimshony *et al*., 2016) with alterations similar to Nelms & Walbot (2019). The samples were thawed then incubated at 65□ for 3 min, spun, then incubated again at 65□ for 3 min then placed on ice. To each sample 0.7 µL of reverse transcription mix (8:2:1:1 of Superscript IV 5X Buffer, 100 mM DTT, RNase Inhibitor, Superscript IV; ThermoFisher Scientific) was added, spun down, then incubated at 42□ for 2 min, 50□ for 15 min, 55□ for 10 min then placed on ice. The samples were pooled by row into 8-strip tubes and excess primers were digested with the addition of 4.6 µL exonuclease I mix (2.5 µL of 10X Exonuclease I Buffer, 2.1 µL Exonuclease I; New England Biolabs) then incubated at 37□ for 20 min, 80□ for 10 min then placed on ice. To each of the pooled samples 44.28 µL (1.8X volume) of pre-warmed RNAClean XP beads was added and mixed well via pipette. The samples were left to incubate at RT for 15 min then placed on a magnetic rack until the liquid became clear. The supernatant was carefully pipetted out, making sure not to disturb the beads, and discarded. The beads were washed twice with 100 µL of freshly prepared 80% ethanol. The ethanol was pipetted out then the beads were left to dry for five min. The RNA was eluted from the beads with 7 µL RNase-free water and incubated for two min at RT then mixed via pipette.

Second strand synthesis was initiated with the addition of 3 µL second strand synthesis mix (2.31 µL Second Strand Reaction dNTP-free Buffer, 0.23 µL 10 mM dNTPs, 0.08 µL DNA ligase, 0.3 µL DNA polymerase I, 0.08 µL RNase H; New England Biolabs) and then incubated at 16□ for 4 h. Samples were further pooled into a single tube and 30 µL Ampure XP beads (Beckman Coulter Life Sciences) with 66 µL bead binding buffer (2.5 M NaCl, 20% PEG 8000; Sigma-Aldrich) (1.2X volume) was added. The sample was incubated for 15 min at RT then washed and dried as described for the RNAClean XP beads above. The RNA was eluted from the beads with 6.4 µL of RNase-free water, left to incubate for 2 min at RT, and mixed via pipette.

*In vitro* transcription was initiated with the addition of 9.6 µL of MegaScript T7 IVT mix (1:1:1:1:1:1 of CTP solution, GTP solution, UTP solution, ATP solution, 10X Reaction Buffer, T7 Enzyme Mix; ThermoFisher Scientific) to the sample then incubated at 37□ overnight. The beads were removed from the sample with a magnetic rack and 28.8 µL (1.8X volume) of pre-warmed RNAClean XP beads (Beckman Coulter Life Sciences) was added then incubated at RT for 15 min then washed and dried as described above. Once dry, 6.5 µL of RNase-free water was added to the beads, incubated for 2 min at RT, and mixed via pipette. The amplified RNA quality and quantity were analyzed with an RNA Pico 6000 chip on an Agilent 2100 BioAnalyzer (Agilent Technologies).

To the samples 1.5 µL of priming mix (9:5:1 of RNase-free water, 10 mM dNTPs, 1M tagged random hexamer primer: 5’-GCCTTGGCACCCGAGAATTCCANNNNNN) was added and incubated at 65□ for 5 min then placed on ice. A second round of reverse transcription was initiated with the addition of 4 µL of reverse transcription mix (4:2:1:1 of First Strand Buffer, 0.1 M DTT, RNaseOUT, SuperScript II; ThermoFisher Scientific) to each sample then incubated at 25□ for 10 min, 42□ for 1 h, and 70□ for 10 min before being placed on ice. For the final PCR, 5.5 µL of sample were added to 21 µL of PCR master mix with Illumina TruSeq Small RNA PCR primer (RP1) and Index Adaptor (RPI “X”) (6.5 µL RNase-free water, 12.5 µL Ultra II Q5 Master Mix, 1 µL of 10 µM RP1, 1 µL of 10 µM RPI “X”). Libraries were amplified with 13 rounds of PCR (98□ for 30 sec, then 13 cycles of 98□ for 10 sec, 65□ for 15 sec, and 72□ for 30 sec and finished with 72□ for 3 min). The final PCR products were purified with 26.5 µL (1.0X volume) of Ampure XP beads (Beckman Coulter Life Sciences) then incubated at RT for 15 min then washed and dried as described above. The cDNA was eluted from the beads with 25 µL RNase-free water and purified again with 25 µL (1.0X volume) of Ampure XP beads (Beckman Coulter Life Sciences) then incubated at RT for 15 min then washed and dried as described above. The final purified libraries were eluted into 10 µL RNase-free water incubated for 2 min at RT and mixed via pipette. The cDNA was then assessed with an Agilent BioAnalyzer High Sensitivity DNA chip.

Two libraries of 96 cells isolated with the BioSorter were sequenced on a HiSeqX and two libraries of 96 cells isolated with the Hana were sequenced on a NovoSeq (Illumina) at Novogene Co. (Sacramento, CA, USA) with paired-end 150 base-pair (bp) reads. Primer sequences can be found in Extended Data Table 1-2. All primers were synthesized by the Stanford Protein and Nucleic Acid Facility (PAN, Stanford University, Stanford, CA, USA). Detailed step-by-step protocols of enzyme RNase-depletion, fixed cell isolation, and library preparation can be found in the Supplementary Materials.

### Read filtering, mapping, and initial processing

Paired-end raw reads were demultiplexed based on cell-specific barcodes (Extended Table 1) using Fastq-Multx (Aronesty, 2013). The UMI sequences from read 1 were added to the read 2 sequence names and then filtered and trimmed with Fastp (parameters: -y -x -3 -f 6) (Chen *et al*., 2018). The clean reads were mapped to the B73 reference genome (AGP v. 4) (Jiao *et al*., 2017) with HiSat2 (Kim *et al*., 2019), and unique molecular identifiers (UMIs) quantified with SAMtools (Li *et al*., 2009) and UMI-tools (Smith *et al*., 2017). Cell cycle heterogeneity has been shown to distort the clustering of cells, thus all cell-cycle genes from Nelms and Walbot (2019) were removed and cells with fewer than 500 UMIs or 200 genes detected were discarded. Genes that were detected in fewer than 3 cells were also discarded.

**Table 1.**
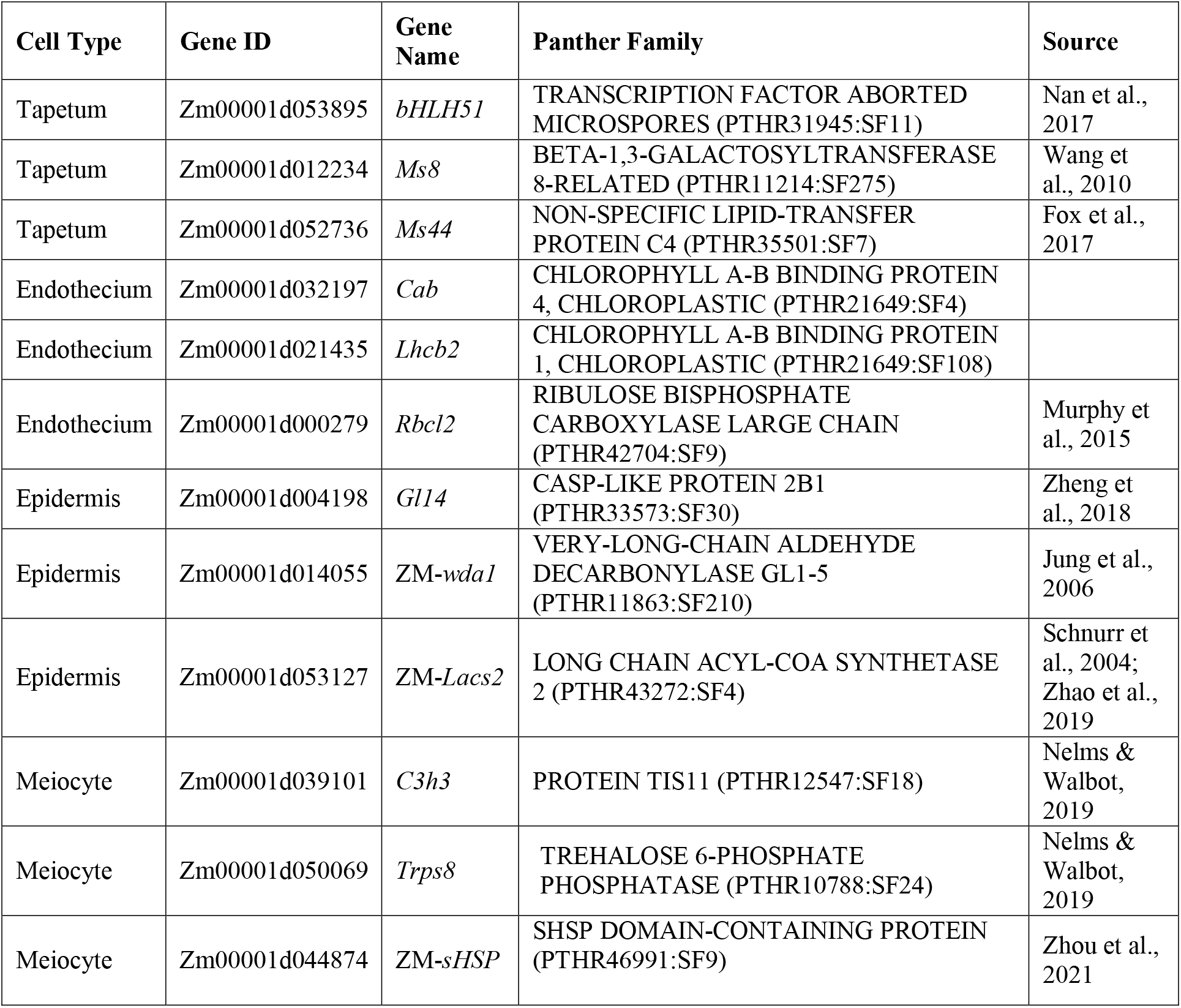
Marker genes and source.

To initially compare our dataset with that of known cell types we assessed the similarity of our data with laser-capture microdissection (LCM) sequencing data of known cell types and whole anthers (Zhou *et al*., 2022), which were also prepared from 2.0 mm W23 maize anthers using the same CEL-Seq2 library preparation. UMIs were normalized into transcripts per million (TPM) and log transformed after adding a pseudocount of 100. We then subtracted the single cell TPMs by the log transformed TPMs of the whole anthers to produce ratio measurements. The LCM data had samples for tapetal, meiocyte, and other somatic (middle layer, endothecium, epidermis) cell types and were similarly processed relative to the whole anther data. We then calculated the cell-to-cell Pearson’s correlations of all our single cells relative to each of the LCM samples.

### Cell clustering and cell type identification

Cell clustering and cell type analyses were performed using Monocle 3 (Cao *et al*., 2019) in R/RStudio (R Core Team, 2013; Team, 2015). The UMI counts were normalized via log and size factor with an added pseudocount of 1 and dimensionality reduced via Principal Component Analysis (PCA) consisting of 10 principal components based on the leveling point of an elbow plot of the percentage of variance explained by ranked principal components. Batch effects were removed with the align_cds function in Monocle. Clusters were determined and visualized with Uniform Manifold Approximation and Projection (UMAP) with a resolution of 0.01 (McInnes *et al*., 2020). Correlation values of each cell with the LCM tapetal, meiocyte, and other somatic cell types were mapped onto the UMAP, as well as the percentages of transcripts from the plastid genome and mitochondrial genome. The meiocyte cluster was manually separated from the endothecium cluster based on the LCM correlation data and meiocyte marker genes; it is likely that Monocle did not separate these clusters due to the scarcity of meiocyte cells despite the clear separation in the UMAP. The other somatic 1 (OS1) cluster was subset and reclustered to identify and separate the epidermis cluster based on putative marker genes of the known biology of the cell type (Table 1).

*De novo* cluster-specific marker genes were identified and ranked using pseudo R_2_ values from the marker_test_res function in Monocle. Co-regulated genes were grouped into modules by using the graph_test function to calculate Moran’s I for each gene then applying the Louvian community analysis with a resolution of 0.01 via the find_gene_modules function. We then plotted the aggregate expression of all genes per module for each UMAP cluster to identify cluster-enriched gene modules. The genes from these cluster-enriched modules were then extracted and analyzed for gene ontology (GO) term enrichment relative to the Maize AGPv.4 reference in AgriGO v2 (Tian *et al*., 2017).

## Data availability

Sequencing data are deposited in the NCBI Gene Expression Omnibus under BioProject PRJNA760550.

## ACKNOWLEDGMENTS

This work was supported by the National Science Foundation awards 1907220 (to DBM) and 17540974 (to B. C. Meyers and V.W.). We thank Ed Buckler for sequencing of the preliminary libraries.

## AUTHOR CONTRIBUTIONS

DBM and BN performed most experiments and designed the study. BN designed and optimized the enzyme RNase-depletion protocol. DBM analyzed the RNA-seq data. DBM wrote the manuscript with input from BN and VW.

## COMPETING INTERESTS

A patent on the enzyme RNase-depletion protocol has been filed by Stanford University with BN as inventor (U.S. Patent Application No. 17/196,681).

## EXTENDED DATA FIGURE LEGENDS

**Extended Data Fig. 1.**
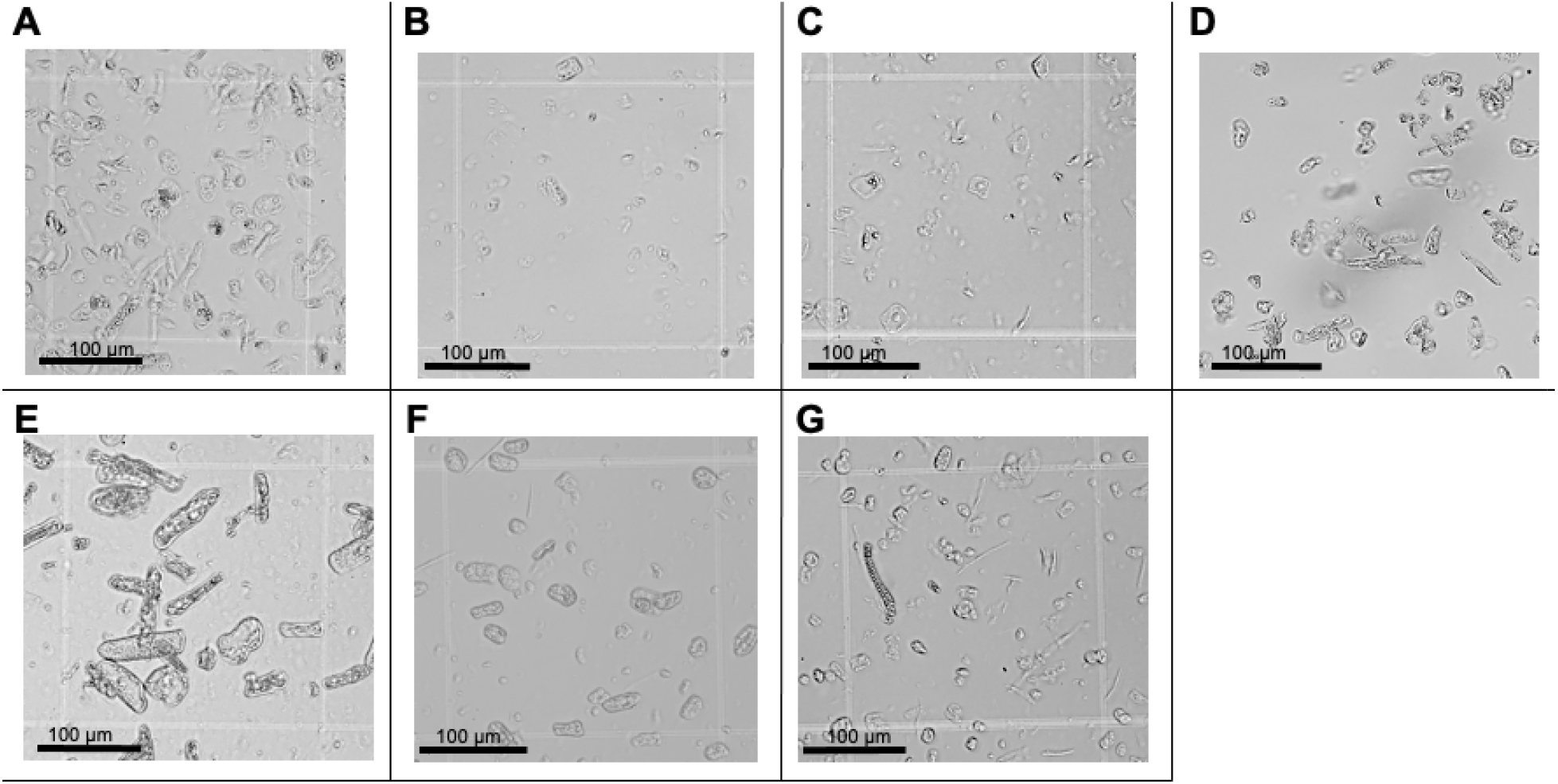
Cell dissociation and morphology via FX-Cell. Fixed (A) maize apical meristem cells, (B) maize leaf cells, (C) maize ear cells, (D) *Amborella trichopoda* leaf cells, (E) *Nymphaea colorata* leaf cells, (F) *Capsella bursa-pastoris* leaf cells, and (G) *Capsella bursa-pastoris* stem cells after digestion and mechanical dissociation.

**Extended Data Fig. 2.**
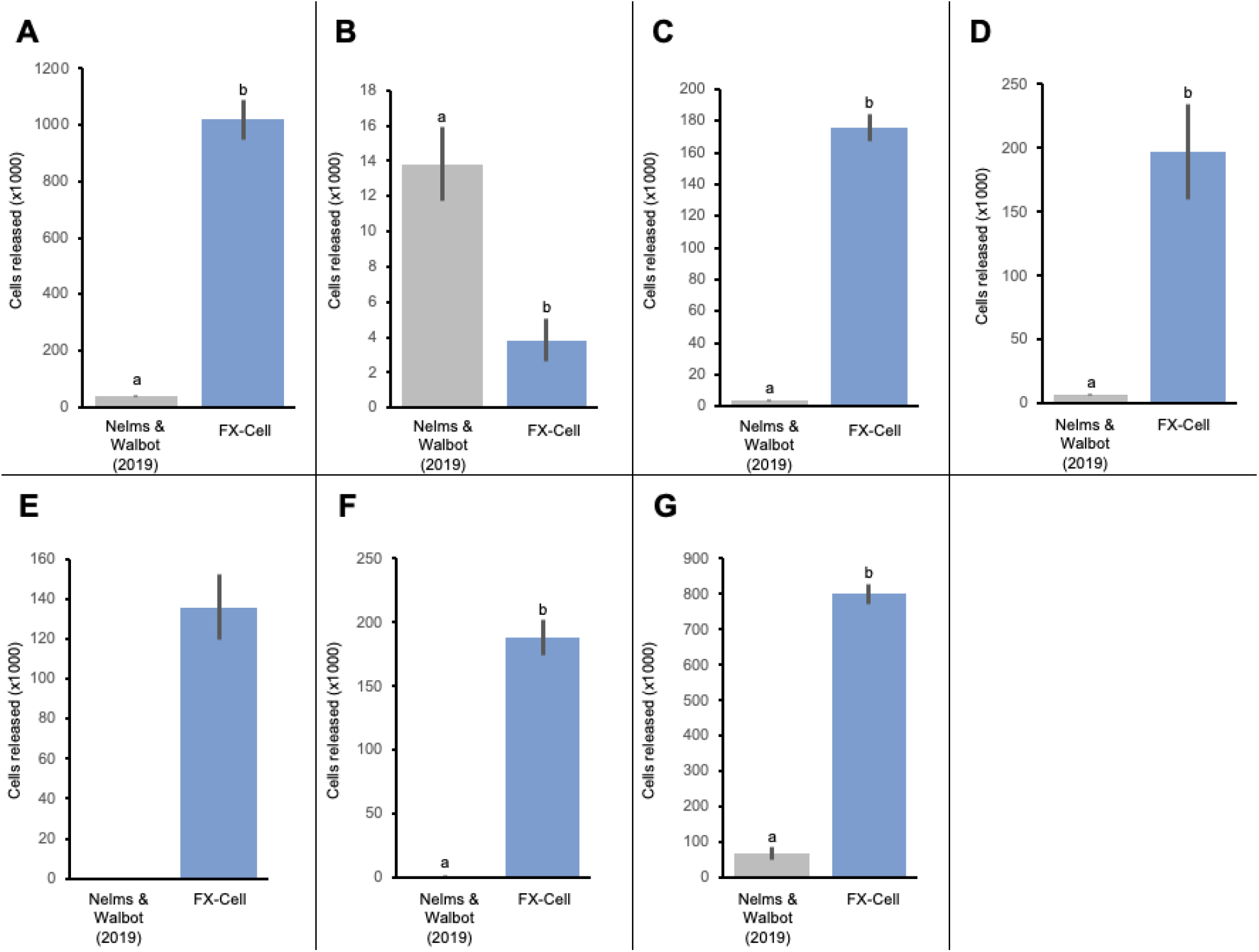
Cellular release of varying plant tissues via protoplasting and FX-Cell. Number of individual cells released using fresh protoplasting (Nelms and Walbot, 2019) or FX-Cell from (A) maize apical meristem tissue, (B) maize leaf tissue, (C) maize ear tissue, (D) *Amborella trichopoda* leaf tissue, (E) *Nymphaea colorata* leaf tissue, (F) *Capsella bursa-pastoris* leaf tissue, and (G) *Capsella bursa-pastoris* stem tissue. Different letters denote statistically significant variation (Student’s *t* test, *P* < 0.05) and error bars represent standard error.

**Extended Data Fig. 3.**
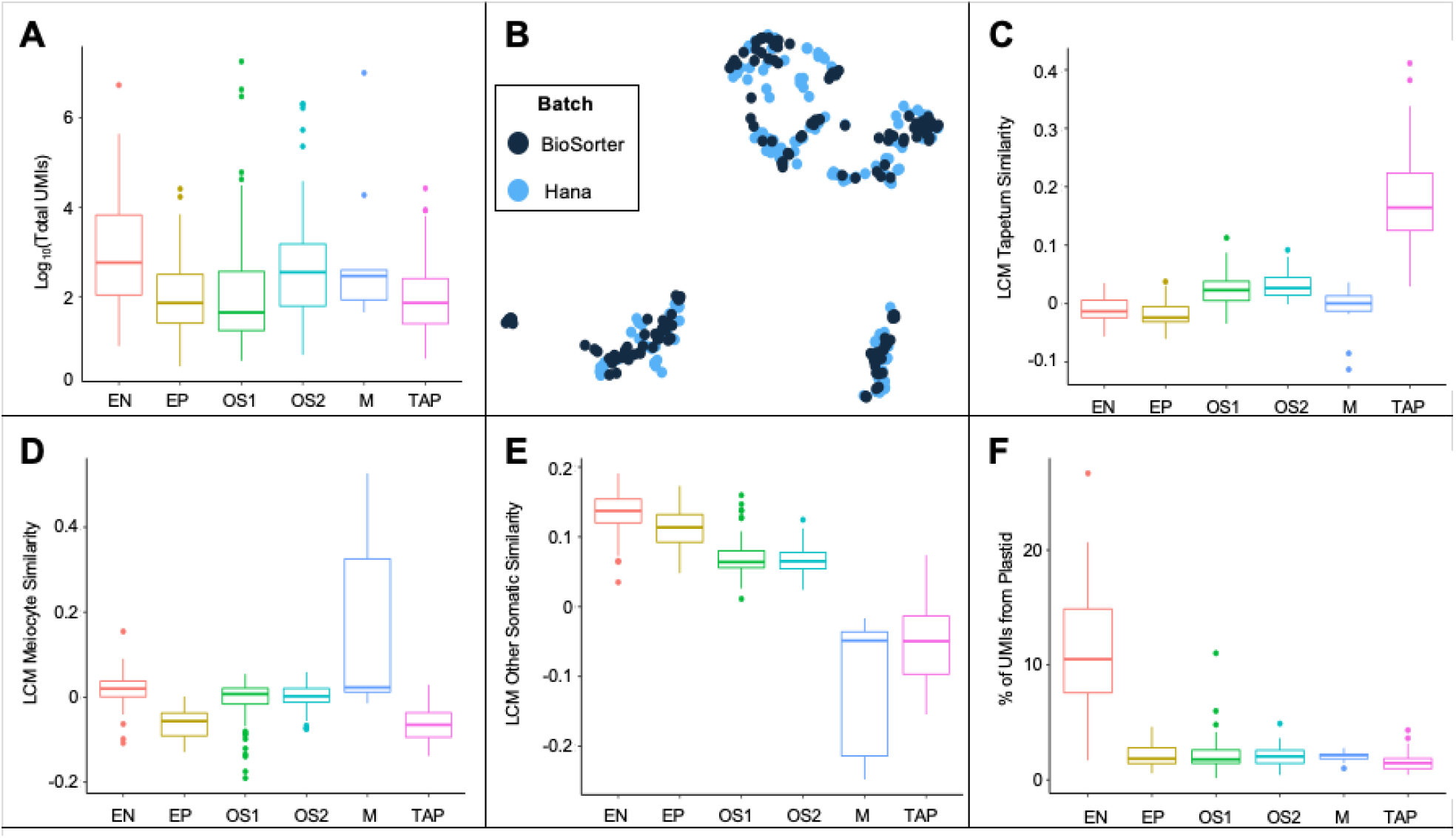
Maize anther scRNA-seq. (A) Box plots of total UMIs per cell by cluster. (B) Cell isolation method (Biosorter vs. Hana) for each cell. (C) Box plots of correlation values with LCM tapetal data for each cell by cluster. (D) Box plots of correlation values with LCM meiocyte data for each cell by cluster. (E) Box plots of correlation values with LCM other somatic cell types (middle layer, endothecium, epidermis) for each cell by cluster. (F) Box plots of percent plastid transcripts for each cell by cluster. The horizontal lines within the box plots represent the median value, the lower and upper bounds of the box plots represent the first and third quartiles, whiskers extend to 1.5x the interquartile range, and all other points are outliers.

**Extended Data Fig. 4.**
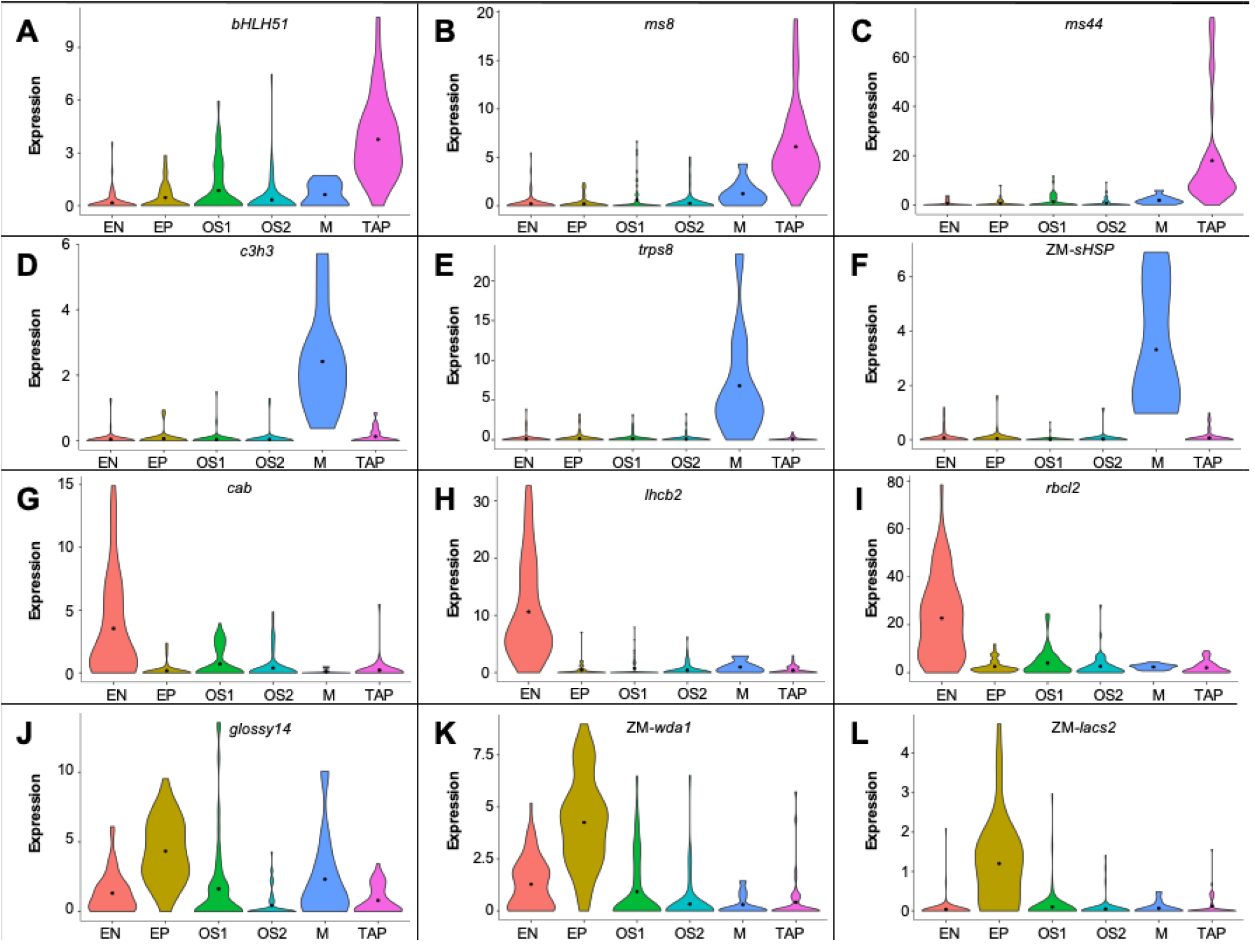
Maize anther scRNA-seq marker gene expression. (A-C) Single-cell RNA-seq violin plots showing expression of tapetal marker genes across the six clusters. (D-F) Single-cell RNA-seq violin plots showing expression of meiocyte marker genes across the six clusters. (G-I) Single-cell RNA-seq violin plots showing expression of endothecium marker genes across the six clusters. (J-L) Single-cell RNA-seq violin plots showing expression of epidermis marker genes across the six clusters.

## EXTENDED DATA TABLES

**Extended Data Table 1.**
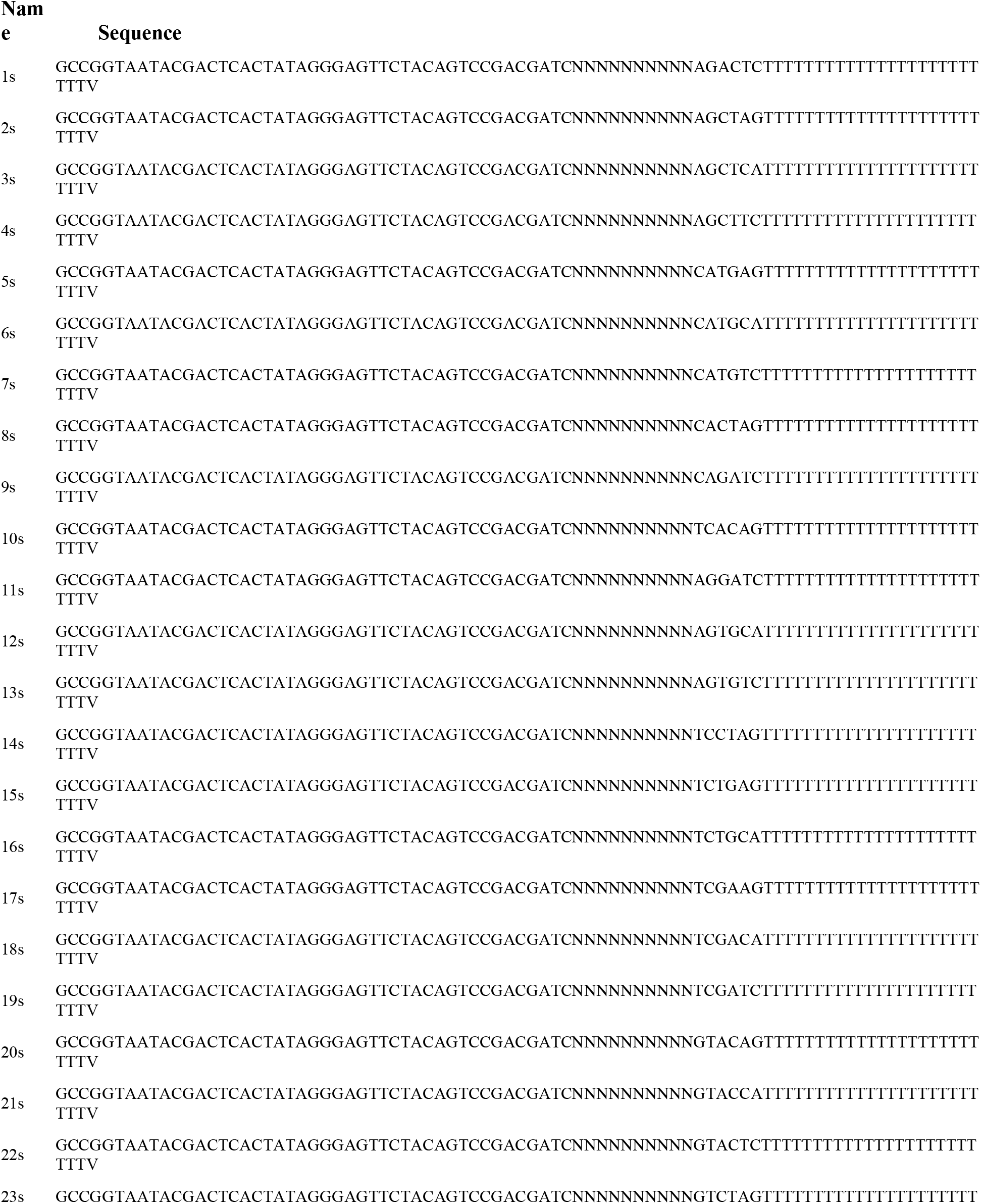
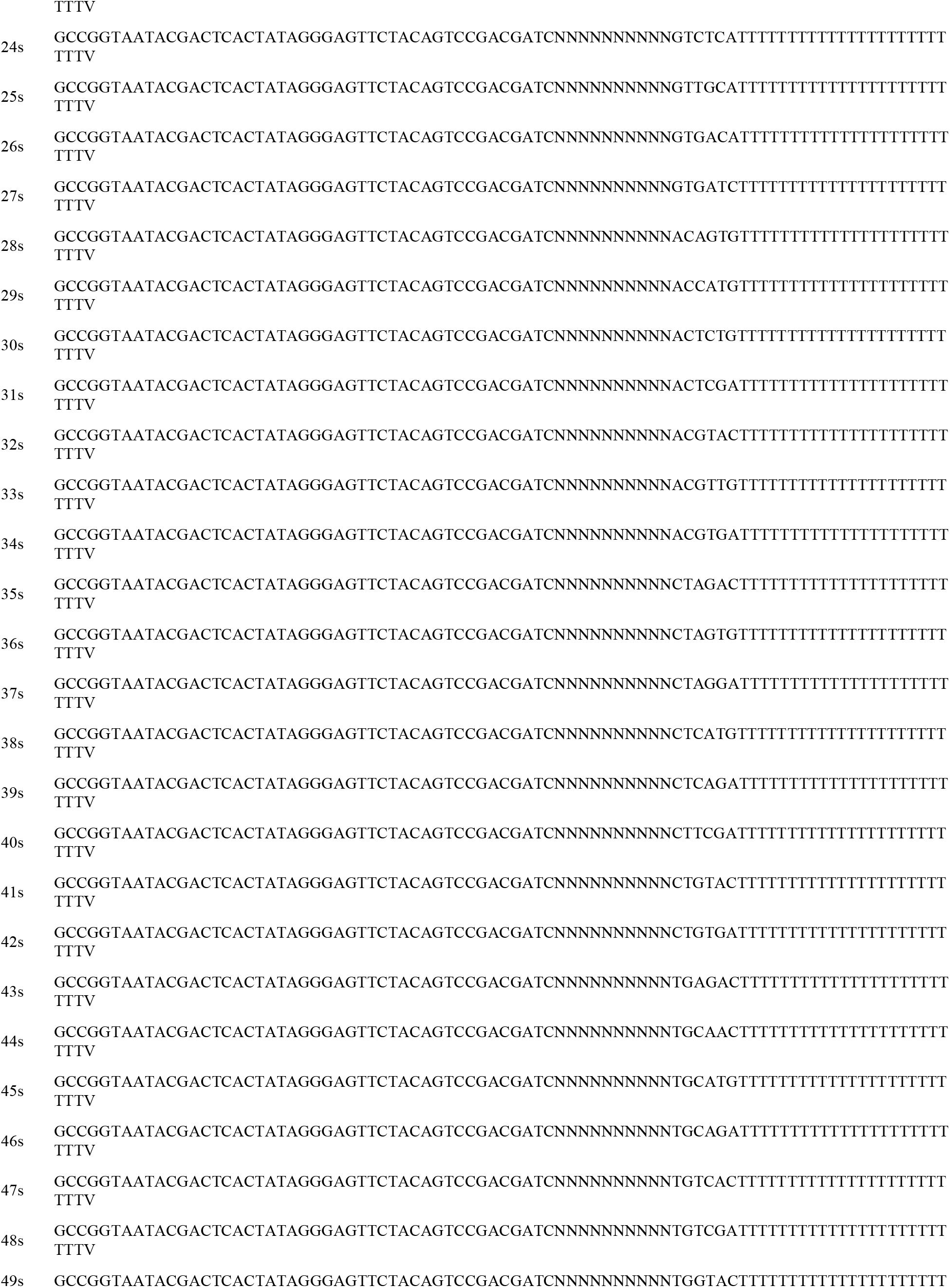
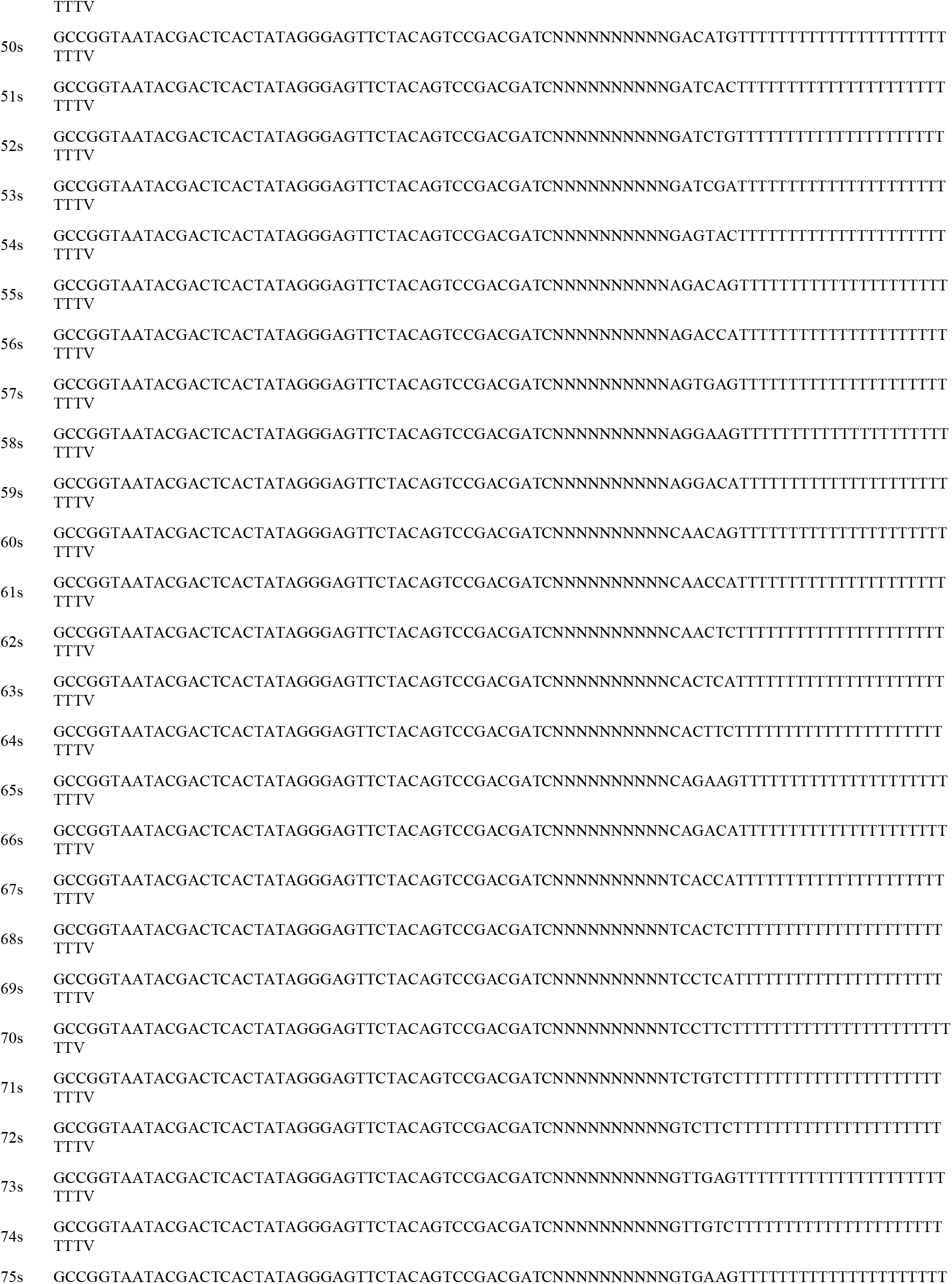
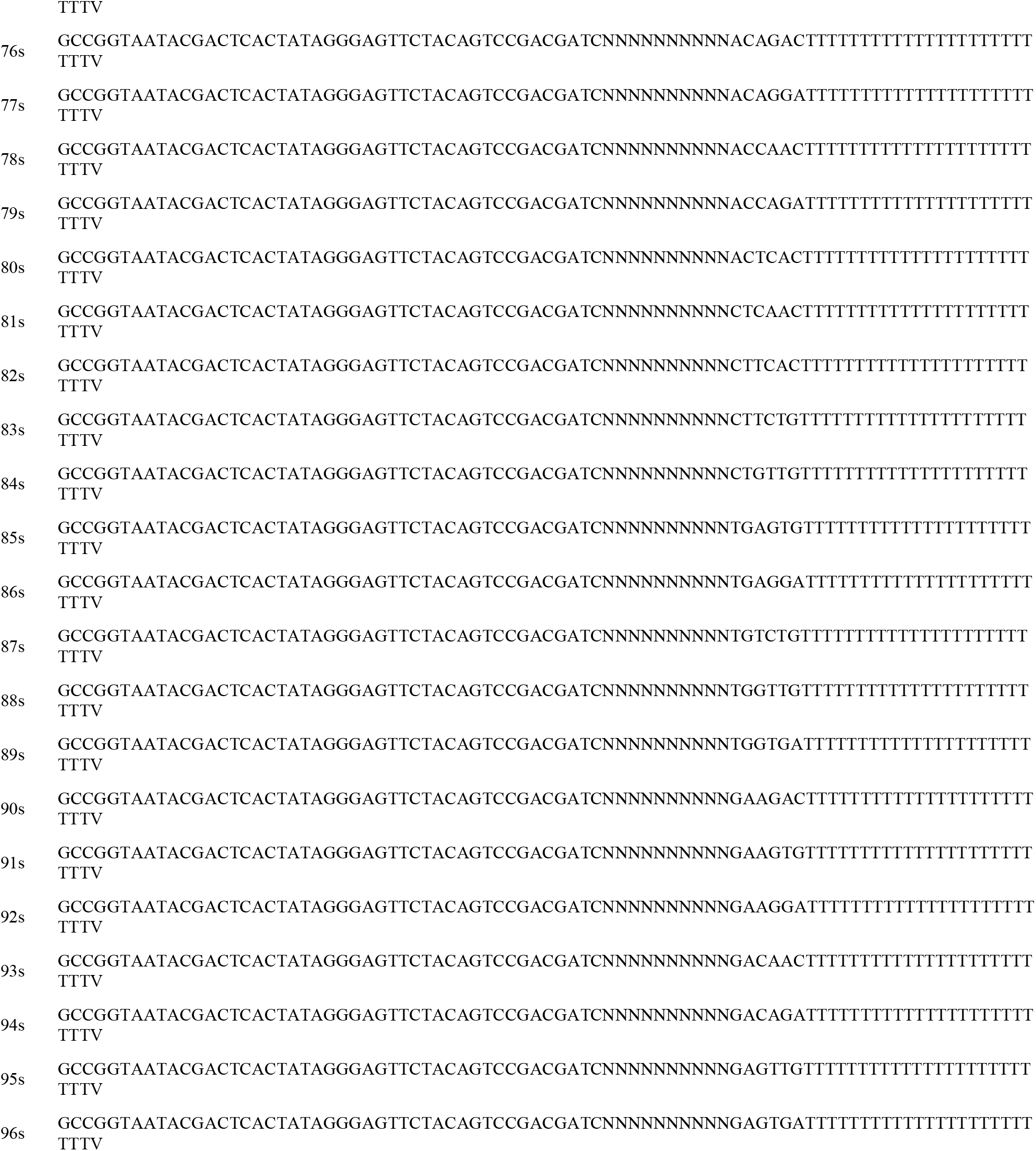
CEL-seq primer sequences (Hashimshony et al 2016) for library construction.

**Extended Data Table 2.**
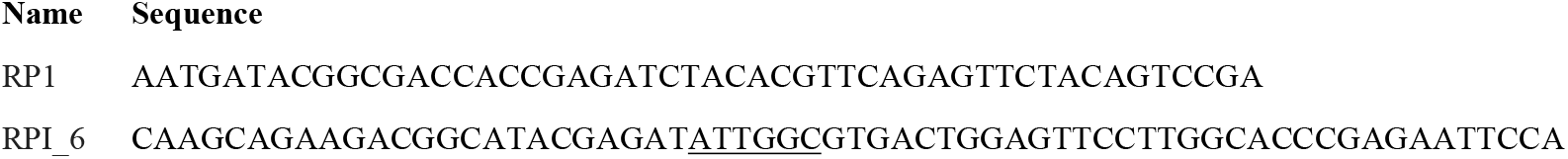
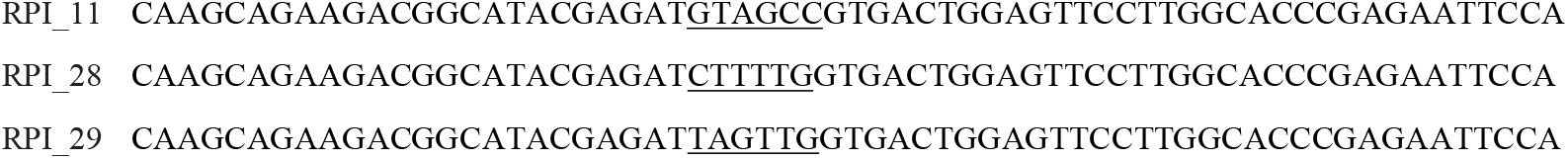
Uniquely indexed RNA PCR primer sequences from Illumina. Barcodes are underlined.

## SUPPLEMENTARY MATERIAL

### DETAILED PROTOCOL

#### Enzyme Purification (Adapted from Kanaya and Uchida 1981)

- Wash 50 mL suspended ω-aminohexyl–agarose beads in water three times.
- Wash beads in 0.1M borax, pH 9.0 three times.
- Separately, dissolve sodium metaperiodate in 6 mL water to a final concentration of 0.2M and add 488 mg GMP.
- Incubate solution at room temperature in the dark for 1 h with gentle mixing.
- Resuspend the washed agarose beads in 0.1M borax, pH 9.0 to a total volume of 36 mL.
- Add 6 mL solution containing oxidized GMP to agarose beads and incubate at room temperature with gentle mixing for 2-4 hours.
- Slowly add 136 mg of solid sodium borohydride to the reaction and gently mix the solution at 4□ for 1 hour with the cap loosened to allow ventilation.
- Wash the coupled GMP beads three times each with 0.1M borax, then water, then 1M sodium chloride.
- Load GMP beads into a FPLC column (ex. Superdex 200 10/300; Cytiva) and store in 1 M sodium chloride at 4□ until ready for enzyme purification.
- Extra beads can be stored in a sealed container in 1M sodium chloride.
- Resuspend enzymes at 10X concentration (12.5% w/v Cellulase-RS and 4% w/v Macerozyme R10) in RNase binding buffer (RBB; 150 mM NaCl, 10 mM citrate, pH 7.0) Load 4 mL of GMP beads in a gravity flow column and equilibrate with RBB at 4□, then pass enzyme mix through column.
- Collect flow-through and run through the pre-equilibrated GMP-agarose FPLC column at 4□ using a peristaltic pump.
- Collect fractions and pool those with A280 absorbance >0.1.
- Concentrate pooled enzymes with an Amicon Ultra-15 Centrifugal Filter Unit, MWCO 30 kDa (MilliporeSigma) at 4□ until a 1:10 dilution of the enzyme blend has an 0.75 absorbance at A280.
- Purified enzymes can be mixed 1:1 with glycerol and stored at -20□.

#### Primer Master Mix (100 µL per well)

- Mix 1687.5 µL 10 mM dNTP solution and 225 µL 10% TritonX-100 in 7713 µL water.
- Dispense 96.25 µL of this solution into each well of a 96-well plate.
- Add 3.75 µL of each 50 µM barcoded oligo[dT] CEL-Seq2 primer into separate wells.
- Vortex well and spin at 400 x *g* for 30 seconds. This is the Primer Plate for aliquoting Primer Master Mix into sample plates and should be stored at -80□.
- When prepping for cell isolation, aliquot 0.8 µL of each Primer Master Mix into wells of a new 96-well plate. Do this transfer in a cold room to reduce evaporation.
- Seal plate with AlumnaSeal and spin at 400 x *g* for 30 seconds.
- Store sample plates at -80□ until ready for cell isolation.

#### Fixed scRNA-Seq Cell Isolation

- Dissect fresh tissue (∼25 mm_2_) into 50 µL ice cold Farmer’s Solution (3:1 100% Ethanol:Glacial acetic acid) in 100 µL 8-strip PCR tubes for two hours. Make sure tissue is submerged.
- Pipette out Farmer’s Solution, add 50 µL ice cold 0.1X Phosphate Buffered Saline (PBS) for five minutes, repeat once.
- Pipette out 0.1X PBS, add 27 µL 20 mM MES and 3 µL 10X purified enzyme, mix well. Make sure tissue is submerged.
- Digest tissue at 50□ for 90 minutes.
- Pipette enzyme solution and tissue up and down ten times to further dissociate cells.
- Transfer enzyme/cell solution to a glass microscope slide with tape on either end to act as a spacer.
- Place a second glass microscope slide on the first and move the two slides back and forth ten times to further dissociate the cells. Confirm that the cells are dissociated with a microscope.
- Add 50 µL 0.1X PBS to each slide to wash and collect the cells into 1 mL 0.1X PBS on ice.
- Add 7 µL SYBR Green I Nucleic Acid Gel Stain (diluted 1:1000) to solution, let incubate on ice for 20 minutes.
- Filter solution through a 40 µm nylon cell strainer into a 50 mL Falcon tube.
- Wash strainer with an additional 2 mL 0.1X PBS.
- Isolate cells into 96-well plates containing 0.8 µL Primer Master Mix with a Hana Single Cell Dispenser or a BioSorter.
- Seal plates with AlumaSeal and spin at 400 x *g* for 30 seconds then store at -80□.

#### CEL-Seq2 Library Preparation (Adapted from Hashimony et al., 2016)

**Keep samples on ice unless otherwise noted**

<u>DAY 1</u>

- Incubate plate with cells and Primer Master Mix at 65□ for 3 minutes, spin at 400 x *g*, incubate again at 65□ for 3 minutes, place on ice.
- To the side of each well (to minimize bubbles) add 0.7 µL of reverse transcription mix (8:2:1:1 of Superscript IV 5X Buffer, 100 mM DTT, RNase Inhibitor, Superscript IV). Spin plate at 400 x *g* for 30 seconds, lightly vortex, then spin again. Incubate plate at 42□ for 2 minutes, 50□for 15 minutes, 55□for 10 minutes then place on ice.
- Pool samples by row into 8-strip tubes, reducing 96 samples to eight.
- To each tube add 4.6 µL exonuclease I mix (2.5 µL of 10X Exonuclease I Buffer, 2.1 µL Exonuclease I).
- Incubate plate at 37□ for 20 minutes, 80□ for 10 minutes then place on ice.
- Add 44.28 µL (1.8X volume) of pre-warmed RNAClean XP beads, mix well, and incubate at room temperature for 15 minutes.
- **Bead Wash**
  - Place sample on magnetic rack until the liquid clears then discard the supernatant, careful not to disturb or pipette up the beads.
  - Add 100 µL of freshly made 80% ethanol, incubate 30 seconds, remove ethanol.
  - Repeat previous step once more.
  - Remove all ethanol and let beads dry for ∼5 minutes.
- Elute with 7 µL RNase-free water and incubate for two minutes at room temperature then mix via pipette.
- Add 3 µL second strand synthesis mix (2.31 µL Second Strand Reaction dNTP-free Buffer, 0.23 µL 10 mM dNTPs, 0.08 µL DNA ligase, 0.3 µL DNA polymerase I, 0.08 µL RNase H).
- Incubate at 16□ for 4 hours.
- Pool the eight samples into a single tube.
- Add 66 µL of bead binding buffer (2.5 M NaCl, 20% PEG 8000) and 30 µL pre-warmed Ampure XP beads (1.2X volume) to pooled samples, mix well, and incubate for 15 minutes at room temperature.
- Bead Wash, elute with 6.4 µL RNase-free water and incubate for two minutes at room temperature then mix via pipette.
- Add 9.6 µL of MegaScript T7 IVT mix (1:1:1:1:1:1 of CTP solution, GTP solution, UTP solution, ATP solution, 10X Reaction Buffer, T7 Enzyme Mix), incubate at 37□ for 13-16 hours.

<u>DAY 2</u>

- Place sample on magnetic rack for 5 minutes and transfer sample without beads into new 100 µL tube.
- Add 28.8 µL (1.8X volume) of pre-warmed RNAClean XP beads and incubate at room temperature for 15 minutes.
- Bead Wash, elute with 6.5 µL of RNase-free water and incubate for two minutes at room temperature then mix via pipette.
- Assess the amplified RNA quality and quantity with an RNA Pico 6000 chip on an Agilent 2100 BioAnalyzer.
  - Expect a primer peak (∼150 bp), but the majority of the sequence distribution should be between 200-1000 bp.
- Samples can be stored at -80□.

<u>DAY 3</u>

- Add 1.5 µL of priming mix (9:5:1 of RNase-free water, 10 mM dNTPs, 1M tagged random hexamer primer) to sample and incubate at 65□ for 5 minutes then place on ice.
- Add 4 µL of reverse transcription mix (4:2:1:1 of First Strand Buffer, 0.1 M DTT, RNaseOUT, SuperScript II) then incubate at 25□ for 10 minutes, 42□ for 1 hour, and 70□ for 10 minutes then place on ice.
- In a new 8-strip tube, add 5.5 µL of sample to 21 µL of final PCR master mix with Illumina TruSeq Small RNA PCR primer (RP1) and Index Adaptor (RPI “X”) (6.5 µL RNase-free water, 12.5 µL Ultra II Q5 Master Mix, 1 µL of 10 µM RP1, 1 µL of 10 µM RPI “X”).
- Optional Amplification Optimization:
  - Transfer 5 µL of sample and PCR mix to new 8-strip tube and add 0.5 µL SYBR Green I Nucleic Acid Gel Stain (diluted 1:5000)
  - Run qRT-PCR with SYBR-sample subset (98□ for 30 seconds, then 25 cycles of98 for 10 seconds, 65□ for 30 seconds, and 72□ for 60 seconds, and finish with 72□ for 10 minutes) to see how many amplification cycles are needed.
  - Based on the qRT amplification plot, the optimal number of cycles is at the transition from exponential phase to non-exponential phase (the point at which the curve starts to plateau).
  - We found 13 cycles to be the optimal number of cycles for all our samples.
- Amplify sample at 98□ for 30 seconds, then X cycles of 98□ for 10 seconds, 65□ for 30 seconds, and 72□ for 60 seconds, and finish with 72□ for 10 minutes.
- Add 26.5 µL, or 20.5 µL if subset removed for optimization, (1.0X volume) of Ampure XP beads to sample and incubate at room temperature for 15 minutes.
- Bead Wash, elute with 25 µL RNase-free water and incubate for two minutes at room temperature then mix via pipette.
- Add 25 µL (1.0X volume) of Ampure XP beads and incubate at room temperature for 15 minutes.
- Bead Wash, elute with 10 µL RNase-free water and incubate for two minutes at room temperature then mix via pipette.
- Assess cDNA product with an Agilent BioAnalyzer High Sensitivity DNA chip.
  - Sequences should be evenly distributed between 200-1000 bp.
  - Expected concentration should be ∼1 ng/µL.
  - Additional size selection with Ampure SPRIselect beads should be applied if a sizeable primer peak (<200 bp) is present.
- Samples can then be stored at -80□.

#### Materials and Reagents

ω -Aminohexyl–Agarose beads (Sigma-Aldrich #A6017)

Guanosine monophosphate (GMP; Santa Cruz Biotechnology #295032)

Borax (Sigma-Aldrich #71997)

Sodium metaperiodate (Chem-Impex #30205)

Sodium borohydride (Sigma-Aldrich #71320)

Sodium chloride (Invitrogen #AM9759)

Amicon Ultra-15 Centrifugal Filter Unit, MWCO 30 kDa (MilliporeSigma #Z717185)

Axygen Low Profile 8-Strip PCR Tubes (Fisher Scientific #14-223-505)

Farmer’s Solution (3:1 100% Ethanol:Glacial Acetic Acid) Poly(ethylene glycol) (Sigma-Aldrich #89510)

Phosphate-buffered saline (PBS; Sigma-Aldrich #P4417)

SYBR Green I Nucleic Acid Gel Stain (Invitrogen #S7563)

Cellulase-RS (Sigma-Aldrich #C0615)

Macerozyme R10 (Sigma-Aldrich #P2401)

40 µm Nylon cell strainer (Corning #07-201-430)

50 mL Falcon Centrifuge tube (Corning #352098)

96-well LoBind PCR plate (Invitrogen #0030129512)

Deoxynucleotide (dNTP) Solution Mix (New England Biolabs #N0447L)

Triton X-100, 10% in water (Sigma-Aldrich #93443)

AlumaSeal CS Sealing Film (Excel Scientific #FCS-25)

Superscript IV Reverse Transcriptase (ThermoFisher Scientific #18090050)

Exonuclease I (New England Biolabs #MO293)

Second Strand Synthesis (dNTP-free) Reaction Buffer (New England Biolabs #B6117S)

DNA Polymerase I (New England Biolabs #M0209)

E. coli DNA Ligase (New England Biolabs #M0205)

RNase H (New England Biolabs #M0297)

Agencourt Ampure XP (Beckman Coulter #A63880)

Agencourt RNAClean XP (Beckman Coulter #A63987)

SPRIselect (Beckman Coulter #B23317)

MegaScript T7 Transcription Kit (Invitrogen #AM1334)

Superscript II Reverse Transcriptase (Invitrogen #18064014)

RNaseOUT Recombinant Ribonuclease Inhibitor (Invitrogen #10777019)

NEBNext Ultra II Q5 Master Mix (New England Biolabs #M0544L)

Illumina TruSeq Small RNA PCR Primer (RP1)

Illumina TruSeq Small RNA PCR Index Adaptors (RPI “X”) RNase-free water (Invitrogen #10977023)

Agilent RNA 6000 Pico Kit (Agilent Technologies #5067-1513)

Agilent DNA High Sensitivity Kit (Agilent Technologies #5067-4626)

#### Instruments

Gravity column (ex. Kontes Flex-Column; Kimble Chase, Vineland, NJ, USA)

FPLC column (ex. Superdex 200 10/300 FPLC; Cytiva, Marlborough, MA, USA)

Peristaltic pump (ex. Minipuls 2; Gilson Medical Electronics, Middleton, WI, USA)

Spectrophotometer (ex. Nanodrop; ThermoFisher Scientific, Waltham, MA, USA)

BioSorter (Union BioMetrica, Holliston, MA, USA) or Hana Single Cell Dispenser (Namocell, Mountain View, CA, USA)

BioAnalyzer 2100 (Agilent Technologies, Santa Clara, CA, USA)

## Notes

### Summary of Updates

Revised text.

